# Intermittent antibiotic treatment of bacterial biofilms favors the rapid evolution of resistance

**DOI:** 10.1101/2022.05.03.490405

**Authors:** Masaru Usui, Yutaka Yoshii, Stanislas Thiriet-Rupert, Jean-Marc Ghigo, Christophe Beloin

## Abstract

The rise of antibiotic resistance in bacterial pathogens is a major health concern and the determinants of this emergence are actively studied. By contrast, although biofilms are an important cause of infections due to their high tolerance to a broad range of antimicrobials, much less is known on the development of antibiotic resistance within the biofilm environment, an issue potentially aggravating the current antibiotic crisis. Here, we compared the occurrence of resistance mutations in pathogenic *Escherichia coli* planktonic and biofilm populations exposed to clinically relevant cycles of lethal treatments with the aminoglycoside antibiotic amikacin. This experimental evolution approach revealed that mutations in *sbmA* and *fusA* are rapidly selected in biofilm but not in planktonic populations. The apparition of these *bona fide* resistance —and not tolerance— mutations is favored by the biofilm preexisting tolerance and high mutation rate. Moreover, we showed that while *fusA* mutations displayed a high fitness cost in planktonic conditions, these mutations were maintained in biofilms, a phenomenon further possibly amplified by the selection of *fimH* mutations favoring biofilm formation itself. Our study therefore provides new insights into the dynamic evolution of antibiotic resistance in biofilms, which could lead to clinically practical antibiotic regimen limiting biofilm-associated infections, while mitigating the emergence of worrisome antibiotic resistance mutations.

## INTRODUCTION

Bacterial infections are a leading cause of morbidity and mortality and the increased propagation of inheritable antibiotic resistance by horizontal gene transfer or the selection of vertically transmitted mutations is a major health concern (Andersson & Hughes, 2017; Murray *et al*, 2022). The use of adaptive laboratory experiment exposing liquid culture planktonic bacteria to sub-inhibitory or progressively increasing concentrations of various antibiotics revealed the rapidity and diversity with which bacteria can acquire inheritable resistance mutations (Jansen *et al*, 2013). By contrast, the use of periodic and short (3 to 8h) lethal antibiotic treatment showed that bacteria first accumulate mutations increasing their tolerance to antibiotics, *i.e.* their ability to survive but not grow under antibiotic pressure, (Fridman *et al*, 2014; Khare & Tavazoie, 2020; Mechler *et al*, 2015; Michiels *et al*, 2016; Sulaiman & Lam, 2020a, b, 2021; Van den Bergh *et al*, 2016b) before to favor, in certain conditions, the emergence of genetic resistance (Levin-Reisman *et al*, 2019; Levin-Reisman *et al*, 2017; Windels *et al*, 2019a). Worryingly, this observation was confirmed in clinical setting in patients treated by a combination of antibiotics for a prolonged *Staphylococcus aureus* bloodstream infection (Liu *et al*, 2020). Although relatively understudied in planktonic conditions compared to resistance (Cohen *et al*, 2013; Van den Bergh *et al*, 2016a; Windels *et al*, 2019b), antibiotic tolerance is a hallmark of bacteria forming surface-attached communities called biofilms (Albert *et al*, 2021; Hall & Mah, 2017; Lebeaux *et al*, 2014). Biofilms indeed display a characteristic high level of tolerance to a broad range of antibiotics that disappears quickly after biofilm dispersion. As a consequence, even when caused by non-resistant bacteria, biofilm-associated infections are difficult to eradicate and regrowth of surviving biofilm bacteria when antibiotic treatment stops is an important cause of therapeutic failures as bacterial infection relapse (Hall & Mah, 2017; Hoiby *et al*, 2015; Lebeaux *et al*., 2014).

Whereas horizontal gene transfer of antibiotic resistance is well-established in biofilms (Chi *et al*, 2007; Ghigo, 2001; Hausner & Wuertz, 1999; Madsen *et al*, 2012; Savage *et al*, 2013; Stalder *et al*, 2020; Strugeon *et al*, 2016), the contribution of biofilm to the emergence of genetically and vertically inheritable antibiotic resistance is less well understood. Yet, biofilms are heterogeneous and diffusion-limited environments that can favor exposure to reduced concentrations of antibiotics (Stewart, 2015). Moreover, the physico-chemical conditions within biofilms induce bacterial stress response and increase mutation rates compared to planktonic conditions (Boles & Singh, 2008; Driffield *et al*, 2008; Jo *et al*, 2022; Martin *et al*, 2016; Steenackers *et al*, 2016; Stewart & Franklin, 2008). All these factors are known to favor bacterial genetic diversification, and, consistently, even in absence of antibiotics, *Escherichia coli* and *S. aureus* biofilms were shown to rapidly evolve resistance to antibiotics (France *et al*, 2019; Koch *et al*, 2014; Tyerman *et al*, 2013). In addition, when exposed to increasing concentrations of daptomycin, *Enterococcus faecalis* biofilms formed in continuous flow conditions evolved resistance mutations correlating with the acquisition of increased biofilm formation capacities (Miller *et al*, 2013). *Acinetobacter baumannii* biofilms treated for 3 days with sub-minimal inhibitory concentration (MIC) of tetracycline or ciprofloxacin developed enhanced survival through enhanced biofilm formation (for tetracycline) or by increasing drug resistance (for ciprofloxacin) (Penesyan *et al*, 2019).

Although the combination of biofilm high antibiotic tolerance with increased genetic diversity is a growing concern, comparison of the evolution of antibiotic resistance in planktonic populations and biofilms led to contradictory results, and no clear evolutionary speed differences could be observed between the two lifestyles. On one hand, *Pseudomonas aeruginosa* colony biofilms continuously subjected to a sub-inhibitory concentration of ciprofloxacin displayed a faster evolution of resistance than planktonic bacteria (Ahmed *et al*, 2018). By contrast, *A. baumannii* biofilms formed on beads and subjected to similar conditions or step-wise increased concentration of ciprofloxacin evolved slower resistance than planktonic populations (Santos-Lopez *et al*, 2019). In the case of *Salmonella* Typhimurium bead biofilms continuously treated with sub-inhibitory concentrations of azithromycin, cefotaxime, and ciprofloxacin (Trampari *et al*, 2021) or *A. baumannii* and *P. aeruginosa* bead biofilms treated with step-wise increased concentration of tobramycin (Scribner *et al*, 2020), no differences were revealed between biofilm and planktonic population evolution rate. In addition, in some of these experiments, higher MICs were reached in evolved planktonic conditions (Ahmed *et al*, 2020; Ahmed *et al*., 2018; Santos-Lopez *et al*., 2019), while in others, no differences in reached MIC could be observed between planktonic and biofilm evolved bacteria (Scribner *et al*., 2020). Hence, whether or not antibiotic-tolerant biofilms constitute a dynamic evolutionary reservoir enabling the emergence of high antibiotic resistance is still unclear.

Here, we performed adaptive evolution experiments to investigate the evolutionary paths toward increased antibiotic resistance in pathogenic *E. coli* planktonic and biofilm populations subjected or not to antibiotic pressure. Considering that the management of chronic biofilm-associated infections often requires the repeated administration of high concentrations of antibiotic (Albert *et al*., 2021; Donelli, 2014; Hoiby *et al*., 2015; Lebeaux *et al*., 2014; Meyer *et al*, 2020), we exposed *E. coli* planktonic and biofilm populations to long (24h) and intermittent treatment with lethal concentrations of the aminoglycoside amikacin. The use of this antibiotic treatment protocol enabled us to show that biofilms evolve enhanced survival to amikacin via a rapid increase of their MIC unlike planktonic populations. By contrast with what was observed with shorter periodic treatments of planktonic populations, neither biofilm nor planktonic bacteria acquired mutations previously associated to antibiotic tolerance, but biofilms accumulated mutations in type 1 fimbriae tip lectin FimH promoting biofilm formation (Yoshida *et al*, 2022) that in turn promoted biofilm-associated tolerance. The evolution of antibiotic resistance in biofilms correlated with the early selection of *sbmA* and *fusA* mutations, increasing resistance to amikacin, which are maintained in the biofilm environment while they displayed a high fitness cost in planktonic conditions. Our study therefore shows that, when biofilms are exposed to clinically relevant intermittent lethal antibiotic treatments, enhanced biofilm tolerance combined with high mutation rates leads directly to the rapid emergence of high level of antibiotic resistance. This suggests that the formation of biofilms in patients subjected to antibiotic treatments constitutes a factor potentially aggravating the foreseen antibiotic crisis. By shedding new light into the evolution of antibiotic resistance in biofilms exposed to clinically relevant antibiotic regimen, the current study could provide insights to improve therapeutic options against biofilm-related bacterial infections.

## RESULTS

### Biofilms evolve enhanced survival to intermittent lethal antibiotic treatment more rapidly than planktonic populations

To investigate the potential emergence of antibiotic resistant mutants in biofilms formed by the pathogenic *E. coli* strain LF82, we used a protocol mimicking the repeated treatments of biofilm-associated infections with amikacin, an antibiotic that is recommended for the treatment of Enterobacterales related infections (Albert *et al*., 2021; Ipekci *et al*, 2014; Pitta *et al*, 2020). We exposed planktonic cultures or biofilms formed on medical-grade silicone coupons to ten cycles of 24h intermittent treatments using 5xMIC (80 µg/mL) and 80xMIC (1280 µg/mL) amikacin, *i.e.* concentrations that are both lethal and superior to the mutation prevention concentration (MPC: 64 µg/mL) (Fig. 1A and 1B). Biofilms treated with such concentrations displayed enhanced survival as compared to planktonic populations (supplementary Fig. S1). At the end of the evolution cycles populations have been propagated for ∼77, ∼200 and ∼36 generations respectively for biofilms treated with 5xMIC, 80xMIC and non-treated, and for ∼324, ∼212 (after 3 cycles, see below) and ∼250 generation respectively for planktonic populations treated with 5xMIC, 80xMIC and non-treated (supplementary Table S1). When submitted to such periodic amikacin treatments, the percentage of biofilm bacteria survival showed an initial drop following the first cycle of 24h treatment and then rapidly increased to reach almost 100% of the population as early as cycle 2 or 3 for the 5xMIC treatment and around 1% of the population for the 80xMIC treatment (Fig. 2A and supplementary Fig. S2A). By contrast, planktonic populations underwent a stronger decrease after the first amikacin treatment and none of the three parallel replicates showed any survivors after 3 cycles with amikacin 80xMIC, while only 0.1% of survival was observed after 7-10 cycles with amikacin 5xMIC (Fig. 2B and supplementary Fig. S2B). These results demonstrated that, whereas 80xMIC treatment showed higher efficiency than 5xMIC on both biofilm and planktonic populations, biofilms were much more resilient to both treatments than planktonic populations, with an increase of the biofilm population size over cycles, which was not observed for treated planktonic populations (supplementary Fig. S3).

**Fig. 1.**
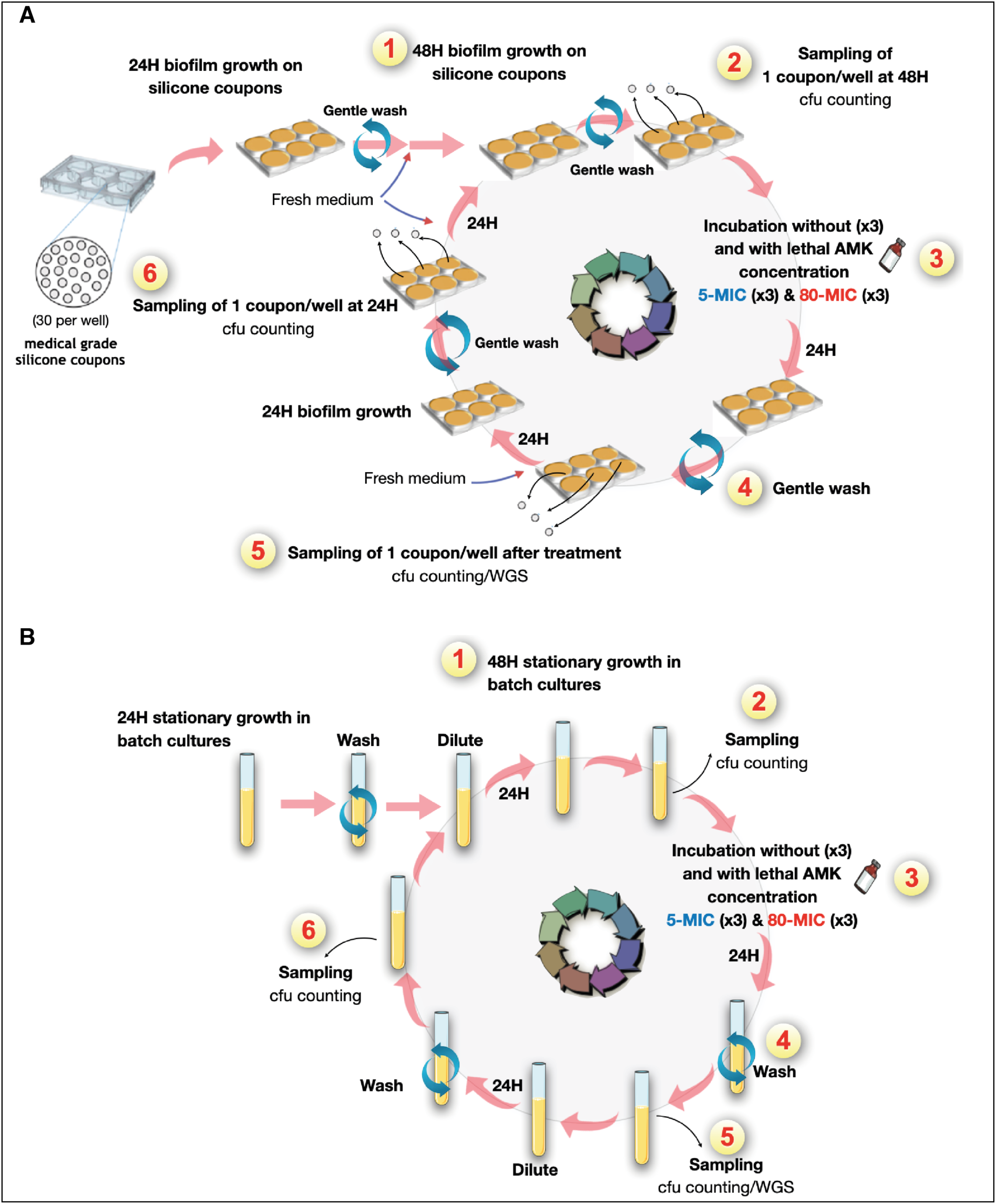
Schematics of experimental evolution in biofilm and planktonic populations under intermittent amikacin exposure with lethal concentrations. (A**) *In vitro* biofilm-model on silicone coupons.** (1) Biofilms of LF82 strains are grown for a total of 48 h at 37°C on silicone coupons in a 6-well plate under static conditions (= cycle 0). (2) One coupon is sampled from each well after removing the exhausted medium and gently washing wells twice with fresh LB medium. (3) Biofilms on silicone coupons are treated with or without amikacin (AMK) at 5- or 80-fold MICs for 24 h at 37°C. (4) Each well is gently washed twice with fresh LB medium after removing the supernatant containing AMK. (5) One coupon is sampled from each well. Next, the survived biofilm cells are incubated in fresh LB medium for 24 h at 37°C. (6) One coupon is collected after washing wells. Then, biofilms are grown for another 24 h at 37°C (= step 1). Steps 2–6 are defined as a series of cycles, and the experiment is performed for 10 cycles. The last cycle was stopped at step 5. Each coupon is collected in a microtube containing LB medium, and the microtubes are vortexed and sonicated to detach biofilm cells from silicone coupons. The bacterial number of collected sample was confirmed by performing CFU counts. Each collected sample is stored at -80°C and characterized later. (B) ***In vitro* planktonic growth.** (1) After washing and diluting aliquots of the 24-h pre-incubated culture of LF82 strains, the aliquots are incubated in fresh LB medium into a test tube for another 24 h at 37°C under shaking condition (= cycle 0). (2) Aliquots of the incubated culture are collected for further analyses. (3) The planktonic culture is treated with or without AMK at 5- or 80-fold MICs for 24 h at 37°C under shaking condition. (4) Aliquots of treated planktonic cells are washed twice with fresh LB medium. (5) Some aliquots of the washed samples are collected. The bacterial number of collected sample was confirmed by performing CFU counts. Next, the remaining aliquots are diluted 100-fold in fresh LB medium and incubated for 24 h at 37°C under shaking condition. Then, the 24-h incubated culture is washed twice with fresh LB medium. (6) Aliquots of washed samples are collected. Next, the remaining aliquots are diluted in fresh LB medium and incubated for another 24 h at 37°C under shaking condition (= step 1). Similar to the biofilm experiment, steps 2–6 are defined as a series of cycles, and the planktonic evolution experiment is performed for 10 cycles. The last cycle was stopped at step 5. Each planktonic sample is stored at -80°C and characterized later, as well as biofilm samples. Also see the Material and Method section.

**Fig. 2.**
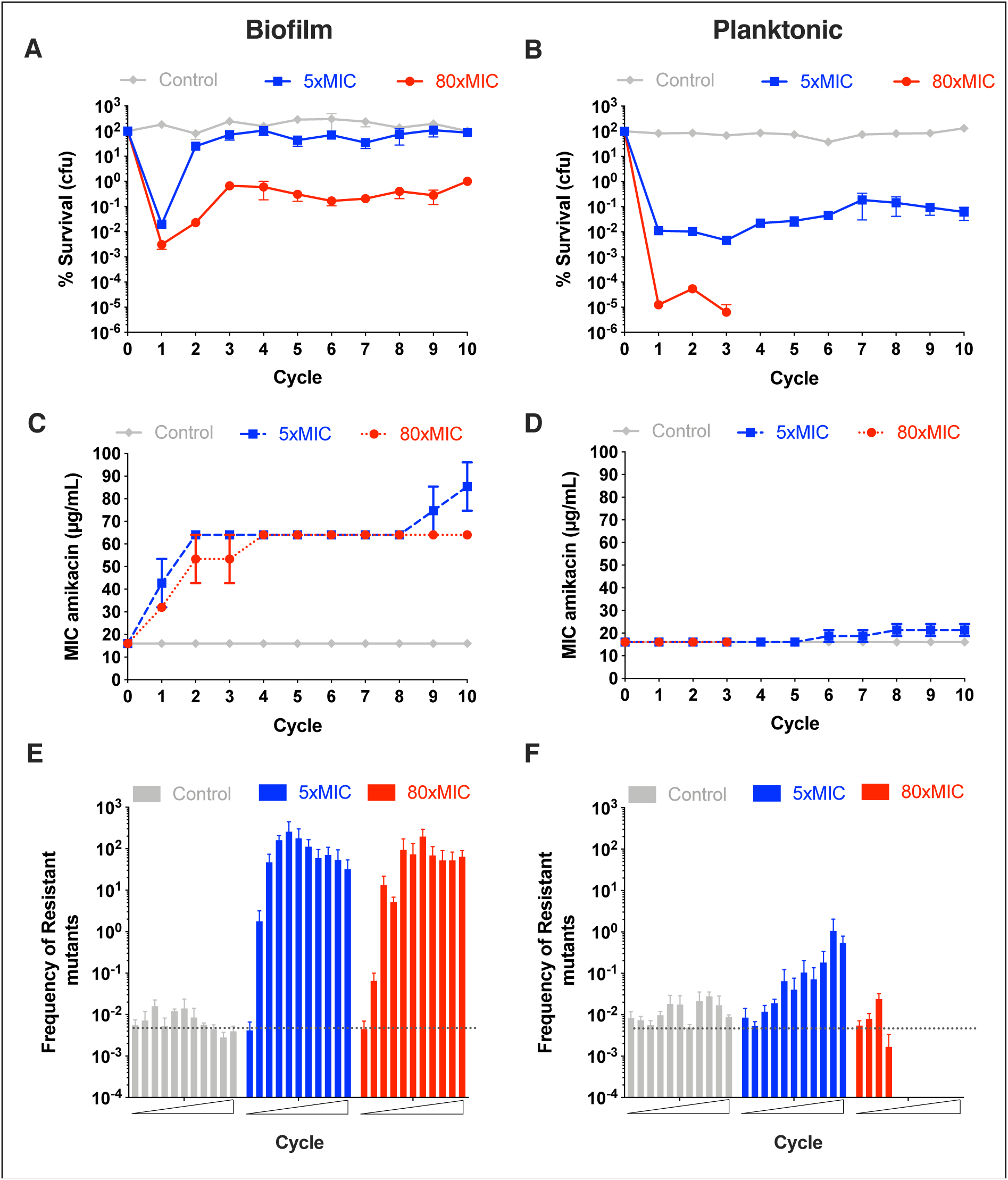
Evolution of *E. coli* survival under lethal antibiotic 24h intermittent treatment of amikacin. (A, C, E) Biofilms and (B, D, F) Planktonic. In (A, B) are represented the percentage of survival after each cycle of evolution at step 5 for biofilms and planktonic as compared to population before each cycle of treatment at step 2 (see supplementary Fig. S1). Percentages of survival were determined using CFU counting. Three independent evolution were performed in parallel for each condition (no treatment = control, 5x amikacin MIC, 80x amikacin MIC). Values represented correspond to the means and standard errors of the mean. In (C, D), the MIC to amikacin was determined for each population sampled at the end of each cycle using the agar dilution method. MICs are represented as the mean and standard errors of the mean of the three independent evolutions for each condition. In (**E, F**), each population sampled at the end of each cycle was plated on LB plate with or without 1x amikacin MIC (or 2xMIC and 4xMIC, see supplementary Fig. S4). Frequency of resistant mutants was calculated as the CFU_1xMIC_/CFU_LB_. For A to E panels, representation of the behavior of individual population is given in supplementary Fig. S2.

### Intermittent antibiotic treatment leads to rapid MIC increase in biofilm but not planktonic populations

To investigate the genetic events underlying the increased survival observed in treated biofilm and planktonic populations, we determined the evolution of their MICs in samples harvested at each treatment cycle. We showed that the evolution of the MICs followed the population survival profiles, with a net and rapid MIC increase in treated biofilms, but only a moderate and late increase in treated planktonic populations (Fig. 2C and 2D and supplementary Fig. S2C and S2D). Consistently, the frequency of clones growing when plated on 1xMIC (Fig. 2E and 2F, supplementary Fig. S2E and S2F), 2xMIC and 4xMIC amikacin (supplementary Fig. S4) increased more rapidly and at lower MIC in treated biofilm samples as compared to treated planktonic ones. Moreover, compared to planktonic samples, the frequency of biofilm clones growing on plates with 2xMIC and 4xMIC was much higher (no planktonic clones grew at 4xMIC) (supplementary Fig. S4). This clearly demonstrated that the evolved clones originating from biofilms displayed a much higher antibiotic resistance than planktonic ones. Finally, we randomly isolated clones from cycle 10 of each evolved population, grew them as planktonic cultures and biofilms, and assessed their survival upon treatment with 5xMIC or 80xMIC amikacin and observed that most of them displayed enhanced survival to 5xMIC or 80xMIC amikacin as compared to their non-evolved parental strain (supplementary Fig. S5). This therefore shows that the higher survival of biofilm clones as compared to planktonic ones correlated with the survival level of the corresponding evolved populations. Moreover, clones isolated from biofilms displayed higher survival rates when grown as biofilms as compared to when grown as planktonic. This suggested that beyond the genetic adaptation of these clones, the biofilms environment, *per se*, plays a critical role in the observed increased survival rates. Therefore, despite being propagated with lower number of generation (supplementary Table S1), biofilms clearly displayed resistance faster and at higher level than planktonic populations.

### The rapid evolution of antibiotic resistance in biofilms correlates with the early selection of mutations in *sbmA, fusA* and *fimH* genes

To identify the genetic bases of the increased survival and MIC levels observed upon intermittent antibiotic treatments, we used whole genome sequencing to sequence cycle 10 end-point biofilm exposed or not to amikacin at 5xMIC or 80xMIC and planktonic populations exposed to amikacin at 5xMIC (Fig. 3 and supplementary Fig. S6 and supplementary Table S2). This analysis revealed mutations in *sbmA* in all treated biofilm populations and 2 out of 3 treated planktonic populations, but not in control non-treated ones. SbmA is an inner-membrane peptide transporter previously associated to increased *E. coli* resistance to aminoglycosides (Jahn *et al*, 2017; Lázár *et al*, 2013; Munck *et al*, 2014) and we determined that most populations with *sbmA* mutations displayed several *sbmA* mutated alleles at various frequencies, indicating potential clonal interference between these different alleles (supplementary Fig. S6 and supplementary Table S2). This was particularly the case in biofilm evolved populations, where 5 out of 6 evolved populations displayed 2 to 5 *sbmA* mutated alleles.

**Fig. 3.**
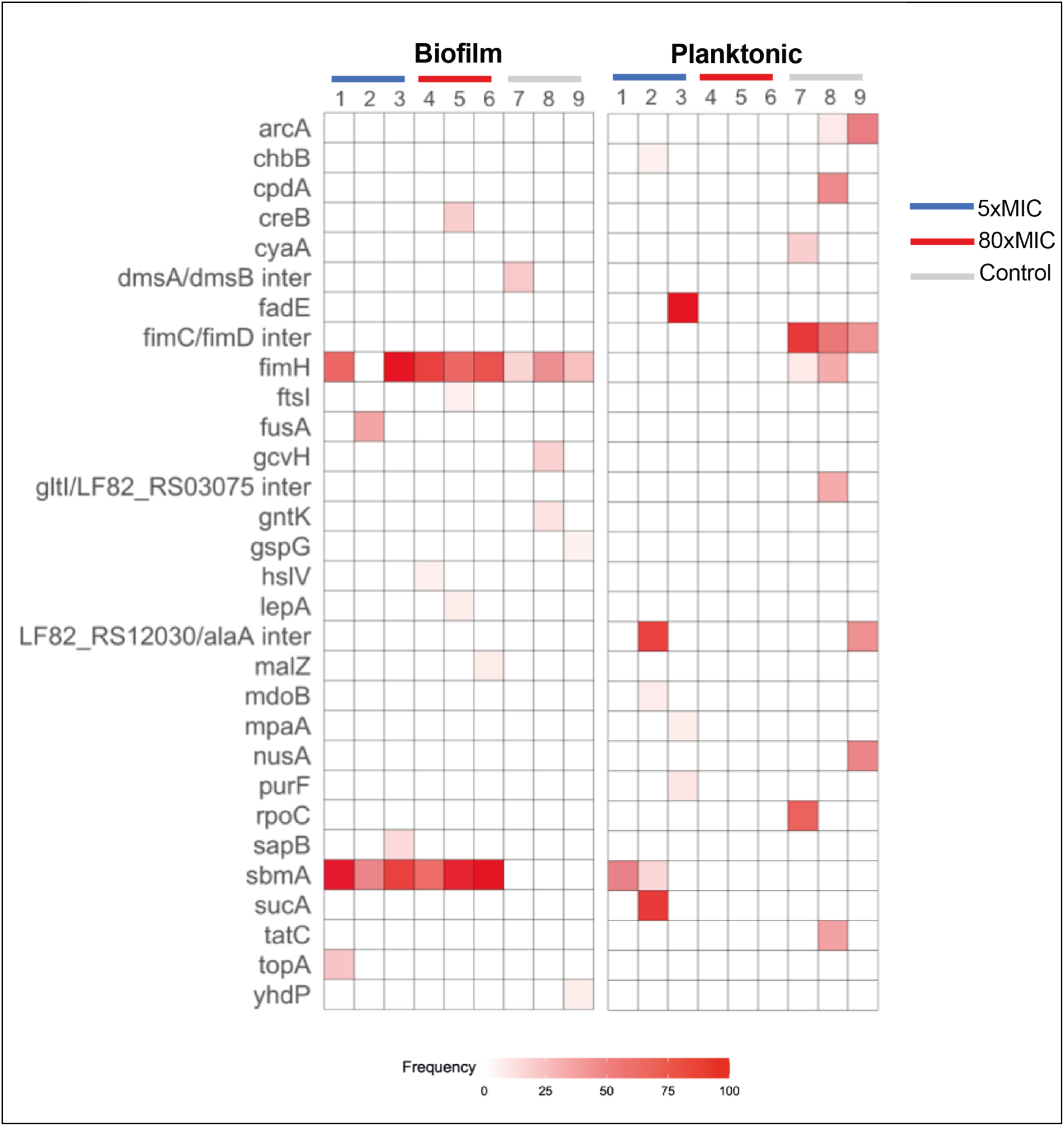
End-point population sequencing reveals a lifestyle associated pattern of mutations after evolution under intermittent antibiotic treatment. Mutations identified by whole-population genome sequencing of control and amikacin treated (5xMIC and 80xMIC) biofilm and planktonic populations of *E. coli*. Three populations per lifestyle and treatment were sequenced after 10 cycles of treatment. Red shading indicates the total frequency of all mutations in each gene within a population at cycle 10 of the experimental evolution. Population 4, 5, 6 from the planktonic lifestyle did not survive after 3 cycles and thus could not be sequenced at cycle 10. We therefore sequenced the population corresponding to the last cycle before their respective extinction. Mutations at higher frequency than 5% are detected by Breseq analysis.

In addition, evolved biofilm population 2 exposed to 5xMIC of amikacin displayed a mutation in *fusA*, which encodes the essential elongation factor G previously associated to increased resistance to aminoglycosides (Ibacache-Quiroga *et al*, 2018; Jahn *et al*., 2017; Lázár *et al*., 2013; Mogre *et al*, 2014; Munck *et al*., 2014). Finally, almost all evolved biofilm populations displayed mutations in the gene coding for the FimH tip-adhesin of type 1 fimbriae (Klemm & Christiansen, 1987). Although these mutations are not directly associated to antibiotic resistance, they might increase biofilm formation and have an impact on survival against antibiotics (see below).

The sequencing of populations at intermediate cycles revealed that the emergence of different *sbmA* alleles in biofilm populations was not only more rapid but also very dynamic and displayed increased diversity of alleles as compared to planktonic populations (Fig. 4 and supplementary Table S3). Moreover, we confirmed the link between presence of *sbmA* mutations and enhanced population resistance since the absence of *sbmA* mutation in evolved planktonic population sample 3 correlated with the absence of MIC increased in this population, which also had the lowest number of colonies growing on amikacin containing plates (supplementary Fig. S2). We confirmed the presence of mutations in *fusA* at intermediate cycles in biofilm populations and showed that, in two populations, *fusA* P610L (sample 1) and *fusA* G604V (sample 4) reached above the 5% frequency detection threshold at cycle 4, while the *fusA* G604V mutation (sample 2) was detected as early as cycle 1 and maintained at high frequency in the population until the end of the experiment (Fig. 4 and supplementary Table S3). At population level, *fusA* and *sbmA* were not mutually exclusive and could coexist in biofilm populations (i.e. in sample 1 and 2) but *fusA* mutations were detected before (sample 2 and 4) or at the same cycle than *sbmA* mutations. No *fusA* mutation could be detected at frequency higher than 5% in evolved planktonic populations.

**Fig. 4.**
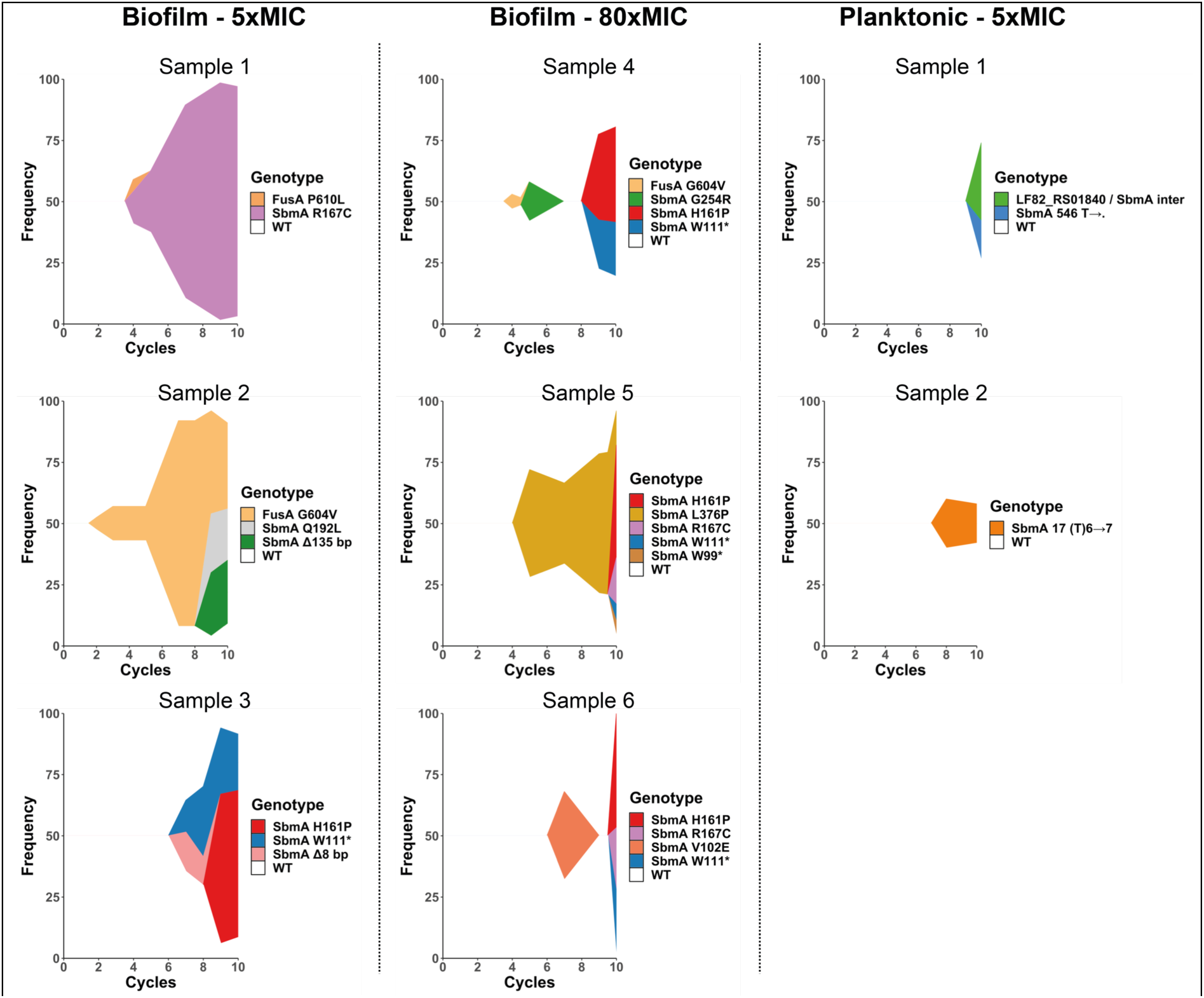
Muller plots showing the dynamics of *sbmA* and *fusA* allele frequencies during the evolution of biofilm and planktonic populations subjected to intermittent amikacin treatment. An independent muller plot showing the dynamic of *sbmA* and *fusA* alleles frequencies over cycles of evolution is shown for each population. Every mutation is displayed with a unique color through all plots. Due to the limit of detection, only mutations with frequency above 5% frequency are shown. Information regarding *sbmA* and *fusA* have been extracted from whole population sequencing presented in supplementary Table S3.

Our analyses therefore showed that in populations treated intermittently with lethal concentration of amikacin, enhanced genetic antibiotic resistance – and not tolerance – is a main driver of the evolution of biofilm survival, mainly due to an early selection of mutations in *sbmA* and *fusA* that could reach high frequency at the end of some of the evolution experiments. By contrast, the weak enhanced resistance of evolved planktonic population emerging in late treatment cycles correlates with rare *sbmA* mutations at a relative lower frequency.

### Dynamics and diversity of mutations leading to amikacin resistance in clones isolated from intermittently treated biofilm and planktonic populations

To get further insights on mutations diversity at clone level, we sequenced the randomly chosen end-point clones analysed for their survival capacity to amikacin (see supplementary Fig. S5) as well as additional randomly chosen clones mostly from cycle 10 and originating from plates with different antibiotic concentration. We also determined their corresponding MICs to amikacin (Fig. 5 and supplementary Table S4).

**Fig. 5.**
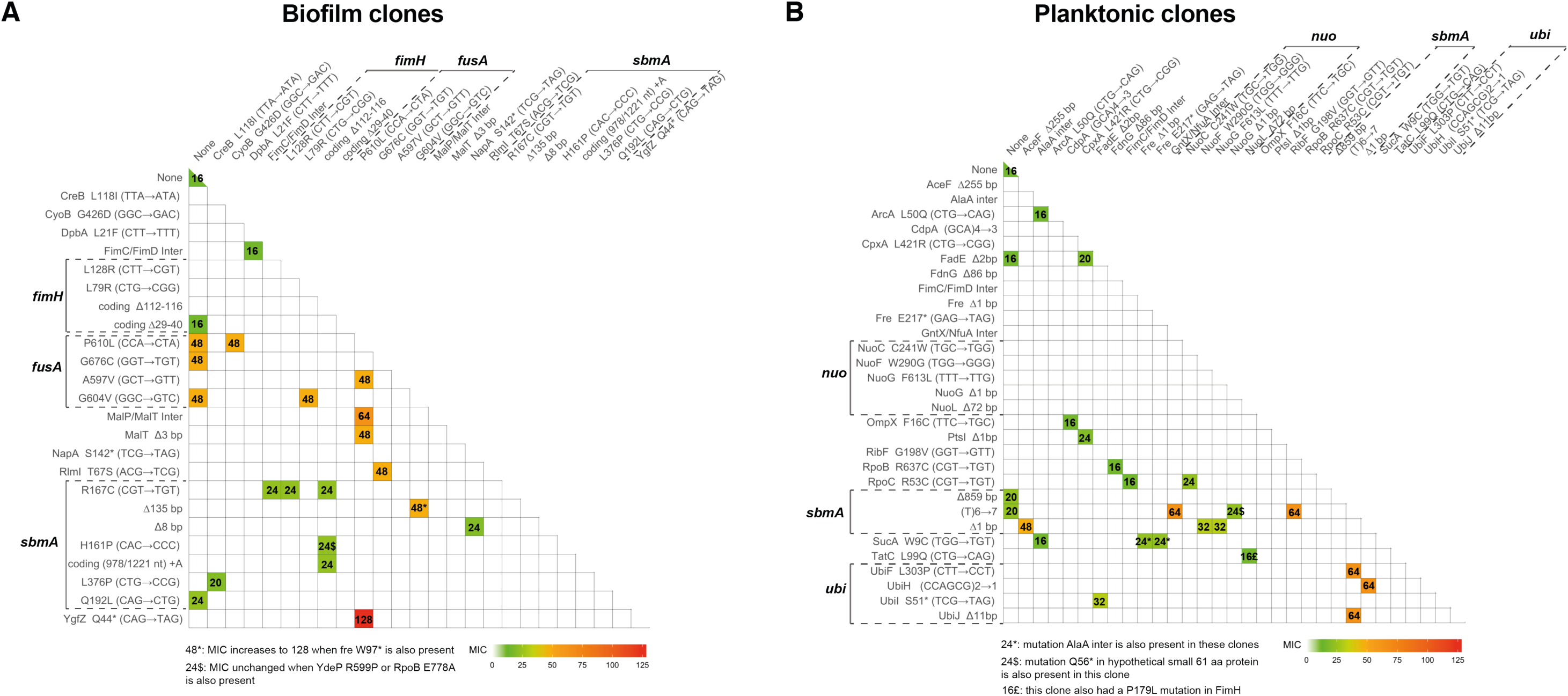
Association of non-synonymous mutations in sequenced evolved clones and their corresponding MICs to amikacin. Non-synonymous mutations identified by whole genome sequencing in clones of *E. coli* coming from amikacin intermittently treated biofilm (A) and planktonic (B) populations. MIC are expressed in µg/mL. MIC of the ancestor strain is 16 µg/mL and is shown in the first top left cell. This information is extracted from clone sequencing data presented in supplementary Table S4.

We first observed that most clones originating from both biofilm and planktonic populations evolved in presence of amikacin exhibited increased MIC compared to the ancestor strain (16 µg/mL) (Fig. 5). In biofilm evolved populations, clones presenting a moderate MIC increase of up to 24 µg/mL of amikacin coded for a loss of function mutation in the *sbmA* gene only, or for mutations not previously linked to antibiotic resistance. This suggested that loss of SbmA activity reduced amikacin efficacy. By contrast, clones with the highest MIC coded for *fusA* mutations, all located in domains IV and V of the elongation factor G protein (supplementary Fig. S7, mutations labeled in red). Whereas the presence of single (G604V, P610L and G676C) or double (A597V P610L) *fusA* mutations led to MICs of 48 µg/mL (Fig. 5), the highest MIC (128 µg/mL) was reached in clones with a mutation in *fusA* and additional loss of function mutations in (i) *sbmA* and in the *fre* NAD(P)H flavin reductase encoding gene or (ii) in *yfgZ,* a gene encoding a protein involved in repair during oxidative stress and Fe-S cluster synthesis (Ote *et al*, 2006; Waller *et al*, 2010) (Fig. 5). The role of YgfZ in Fe-S cluster synthesis could explain a link with aminoglycoside resistance (Ezraty *et al*, 2013). YgfZ is also associated to tRNA modification since a *ygfZ* null mutant has lower levels of several modified nucleotides in tRNA (Ote *et al*., 2006), thus possibly linking increasing of mistranslation with enhanced resistance to some antibiotics (Babosan *et al*, 2022; Witzky *et al*, 2019). Interestingly, whole genome sequencing and Sanger sequencing of the *sbmA* and *fusA* genes in clones from intermediate cycles indicated that the presence of *fusA* mutations could be detected at early cycles of biofilm evolution, suggesting that there were probably already present at low frequencies in the original populations (supplementary Table S4 and S5).

Analysis of clones isolated from evolved planktonic populations revealed, as for biofilm evolved clones, the presence of *sbmA* mutations (Fig. 5). However, except for *sbmA*, the MIC increase of planktonic evolved clones correlated with a higher variety of mutations as compared to the ones found in clones from evolved biofilm populations. These other mutations were nonetheless never detected at frequency higher than 5% in any of the sequenced populations (supplementary Table S3). Moderate MIC increase (20 to 24 µg/mL) in evolved planktonic clones were associated with mutations in *sbmA,* or in *cpxA*, the histidine kinase component of the *cpx* system known to be associated with increase resistance to aminoglycosides through regulation of the expression of porins and efflux pumps (Masi *et al*, 2020). Mutations associated with higher MICs corresponded, in combination or not with *sbmA* alleles, to mutations in genes linked to aminoglycosides resistance via their capacity to modulate the proton motive force (PMF), notably genes encoding components of the Ubi complex responsible for the biosynthesis of ubiquinone (*ubiF, ubiH, ubiI, ubiJ*) (Aussel *et al*, 2014; Muir *et al*, 1981; Xia *et al*, 2017) or genes belonging to NADH-quinone oxidoreductase Nuo (*nuoF*, *nuoG*) (El’Garch *et al*, 2007; Wistrand-Yuen *et al*, 2018). We also identified a clone associating a *sbmA* mutation with a mutation in *aceF*, encoding a component of the pyruvate dehydrogenase complex potentially modulating the efficacy of aminoglycosides through modification of the PMF (Nikaido, 2019; Su *et al*, 2018), and with a mutation in *ribF* involved in riboflavin to FAD transformation that indirectly fuels electron transport chains. A mutation in the intergenic region between *gntX*, encoding a protein necessary for the use of DNA as a carbon source, and *nfuA*, encoding a Fe-S biogenesis protein also enhanced the MIC of a *sbmA* loss of function mutant. NfuA is known to help maturation of NuoG possibly explaining its potential link with resistance to amikacin (Py *et al*, 2012).

Comparison of the MIC of the different clones isolated from biofilm vs planktonic evolution indicated that the MIC of biofilm clones was higher than the one of planktonic (supplementary Table S4, S5 and S6 and supplementary Fig. S8). This supported our previous observation that evolution in biofilms generated clones with higher resistance than evolution in planktonic conditions.

These results show that, at the clone level, biofilm lifestyle allowed the convergent selection of mutations in *sbmA* and *fusA*, although a diversity of mutations in *sbmA* was observed suggesting clonal interference. With the exception of mutations in *sbmA*, the pattern of selected mutations seemed different between biofilm and planktonic evolved populations. Planktonic lifestyle selected clones with mutations in a more diverse set of genes that globally had lower MIC than in biofilms.

### Despite *fusA* mutation fitness cost, the high biofilm mutation rate favors the emergence of amikacin resistance in biofilms

The absence of detectable *fusA* mutants seemed to be one characteristic of evolved planktonic populations. To detect the potential presence of *fusA* mutations at very low frequency that have not been selected and maintained in planktonic populations glycerol stocks of evolved planktonic populations were grown overnight and concentrated 10 times before being plated on 2x amikacin MIC plates. Among the clones growing on these plates we could actually detect a high diversity of mutations in *fusA*, even in the early evolution cycles (supplementary Fig. S7 and supplementary Table S6), confirming the presence of such mutants at very low frequencies at early cycles and their absence of selection. To investigate why *fusA* mutations were selected in biofilm but not planktonic populations, we first determined *sbmA* and *fusA* mutant growth capacity as well as their fitness cost in planktonic conditions. Only P610L *fusA* mutants displayed a relatively lower doubling time (supplementary Fig. S9). However, all three identified *fusA* mutants exhibited a higher fitness cost than the *sbmA* mutants in absence of antibiotics, and this cost was actually still strongly present when the *fusA* mutants were regrown after amikacin treatment (Fig. 6). This suggested that *fusA* mutants could have been counter-selected during intermittent planktonic growth between treatments, while being protected from such counter-selection and maintained in biofilm populations.

**Fig. 6.**
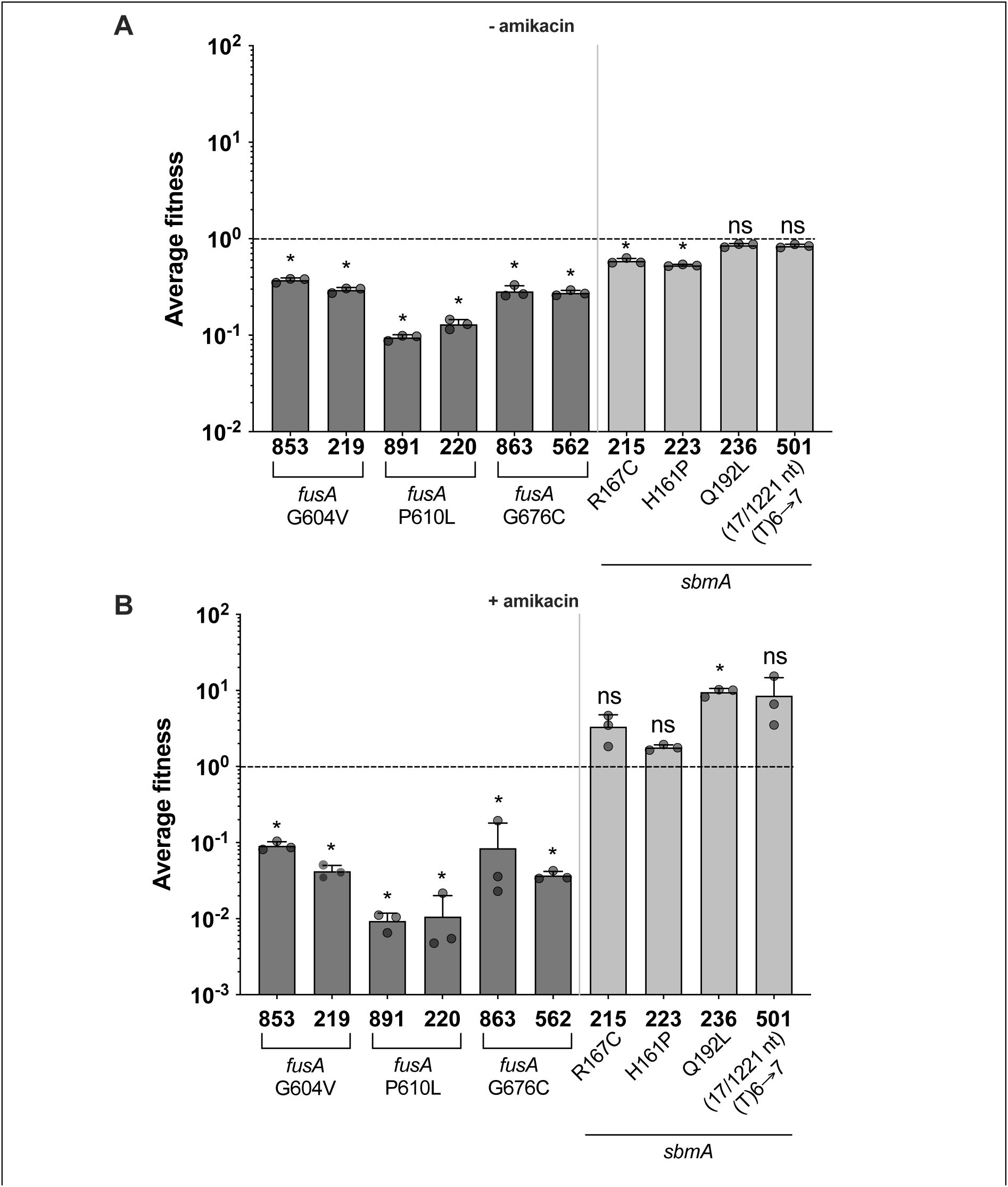
Relative fitness costs of evolved clones containing *fusA* and *sbmA* mutations. Each evolved clone tagged with mars was competed with wildtype tagged with GFP at a 1/1 ratio in the absence (A) and presence (B) of 5x amikacin MIC. In order to match the evolution protocol, in (A) wt/mutants LB overnight cultures were mixed, diluted to OD 0.02 and grown in LB for 24h before flow cytometer analysis. In (B) wt/mutants LB overnight cultures were mixed, diluted to OD 2, treated for 24h by 5x amikacin MIC, washed, diluted 1/100 and regrow in LB for 24h before flow cytometer analysis. The fitness index of the comparison of wildtype tagged with mars to wildtype tagged with GFP is taken as 1 (dotted line). Detailed information on the different analysed clones can be found in supplementary Table S4. The data are represented as the mean ± SD. Statistics correspond to unpaired two-tailed t test with Welch’s correction comparing each condition to wildtype/wildtype fitness. ns, no significance; *, p<0.05; **, p<0.01; ***, p<0.001.

Alternatively, *fusA* mutations could have accumulated at higher frequency in stressful and mutation prone biofilm environment (Beloin *et al*, 2004; Steenackers *et al*., 2016). Consistently, we showed that the mutation rate in the biofilms formed in our experimental evolution set-up was higher than in planktonic conditions (supplementary Fig. S10), therefore providing enhanced opportunity for genetic diversification and the rapid apparition and selection of resistance mutations.

### Mutations enhancing biofilm formation also contribute to enhanced biofilm survival

We recently showed that mutations enhancing adhesion to abiotic surfaces are rapidly selected in the type 1 fimbriae tip lectin FimH during biofilm formation (Yoshida *et al*., 2022). Considering the increased intrinsic tolerance of biofilm to antimicrobials, we wondered whether the high frequency of mutations identified in *fimH* in our evolved biofilm population —all located in the FimH lectin domain (supplementary Fig. S11)— could also contribute to increase biofilm survival (Fig. 3). To evaluate the impact of these mutations on biofilm formation capacity as well as on survival in presence of high amikacin concentration, we tested two of these *fimH* mutations and showed that they indeed promoted both enhanced capacities to form biofilms (Fig. 7A and 7B) as well as increased survival when exposed to 80xMIC of amikacin (Fig. 7C). The differential accumulation and selection of *fimH* mutations in biofilm *vs* planktonic evolution therefore also likely contributed to the rapid and increased capacity of evolved biofilms to sustain high lethal antibiotic concentration.

**Fig. 7.**
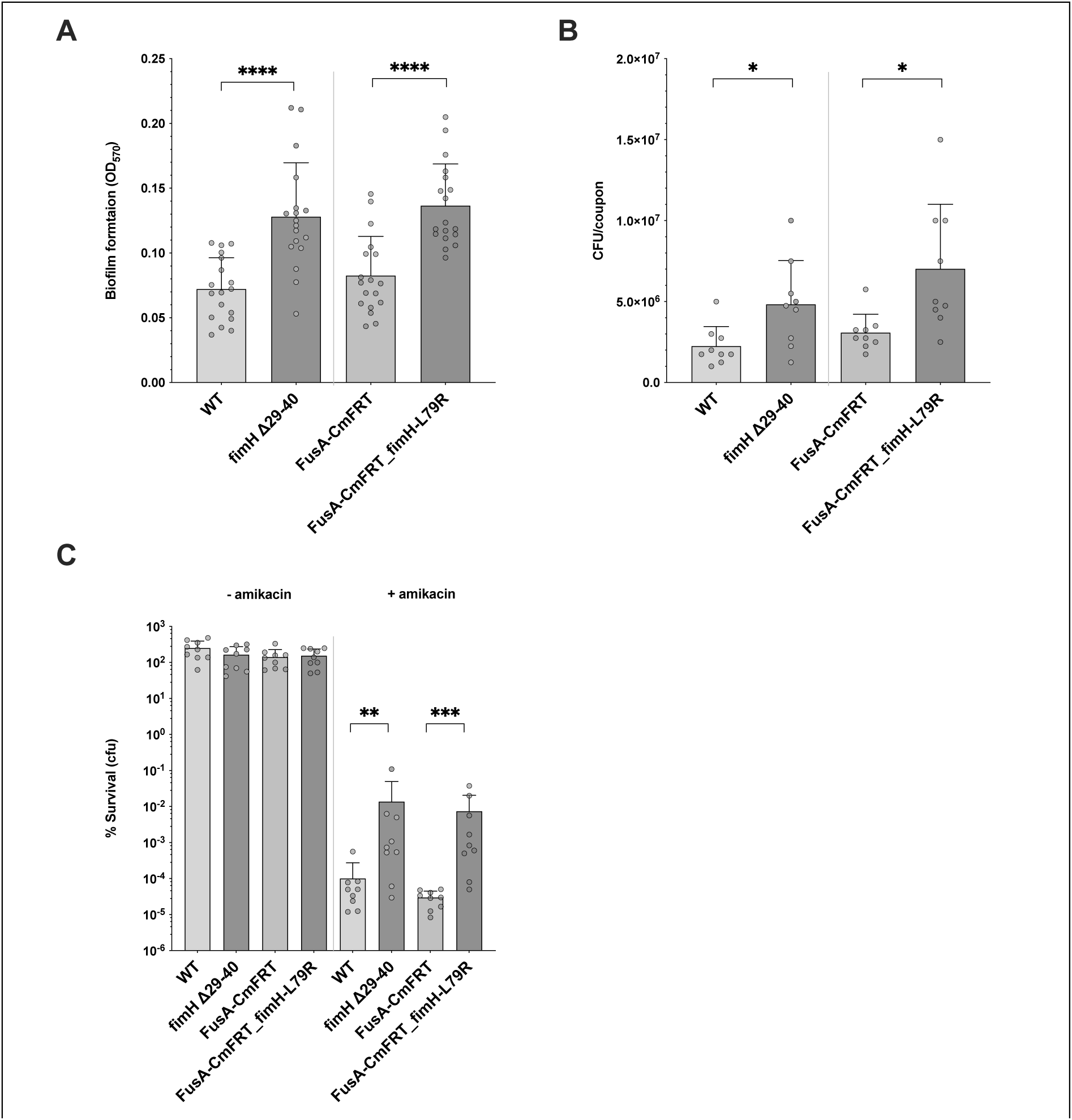
*fimH* mutants display higher capacity to form biofilms and enhanced tolerance towards a lethal concentration of amikacin. (**A** and **B**) Biofilm amounts (**A**) and colony forming units (**B**) of *fimH* mutants formed on silicone coupons for 24 h. (**C**) The survival rate of biofilm cells of *fimH* mutants on silicone coupons after the treatment with 80xMIC of amikacin. The data are represented as the mean ± SD. Statistics correspond to unpaired two tailed t test with Welch’s correction comparing each *fimH* mutant to WT or FusA-CmFRT strains. * p<0.05; ** p<0.01; *** p<0.001, **** p<0.0001. In C, statistics were performed on log-transformed % survival. The clone 239 possesses a unique mutation *fimH* Δ29-40 (see supplementary Table S4). The clone with the mutation *fimH* L79R was created after deconstruction of the *fusA* G604V mutation from the clone 219 that was replaced by a wt version of *fusA* (FusA-CmFRT) (see Material and Methods). This clone also codes for a synonymous mutation I255I in the gene *LF82_RS22000*.

## DISCUSSION

The high but transient tolerance of biofilms to antibiotics and host immune defenses is a well-recognized cause of bacterial chronic infections (Bjarnsholt, 2013; Del Pozo, 2018; Lebeaux *et al*, 2013). There is, however, considerably lower awareness of the risk of emergence of genetically inheritable antibiotic resistance in bacterial biofilms. In this study, we used adaptive experimental evolution and showed that biofilms exposed to periodic treatment with lethal doses of the aminoglycoside antibiotic amikacin, spaced with periods allowing biofilm regrowth, developed antibiotic resistance much faster and at higher level than planktonic populations.

Antibiotic resistance selected through vertical evolution, whereby genetic mutations are heritable and can propagate in the population under antibiotic selection pressure, has been described in many clinical isolates of different bacterial pathogenic species. Whether these mutations arose in patients during or between antibiotic treatments is unclear, especially because such mutations are predicted to occur essentially during long-term or repeated antibiotic treatments (Blanquart, 2019). Although these regimens are characteristic of the clinical management of chronic infections that are often associated with the presence of pathogenic biofilms, few studies established the link between antibiotic treatment of biofilms and the emergence of *de novo* resistance. The most documented cases correspond to the longitudinal analyses of *P. aeruginosa*, *S. aureus,* and *Burkholderia* sp. clones isolated from cystic fibrosis patients subjected to lifelong antibiotic treatments (Goerke & Wolz, 2010; Lieberman *et al*, 2011; Oliver *et al*, 2000; Prickett *et al*, 2017; Rouard *et al*, 2018; Sommer *et al*, 2020; Viberg *et al*, 2017). The emergence of vancomycin, rifampicin or daptomycin resistance has also been reported during *S. aureus* endocarditis or catheter-related bacteremia treatments (Giulieri *et al*, 2018; Kullar *et al*, 2013; Mwangi *et al*, 2007; Riedel *et al*, 2008). Moreover, within-patients antibiotic resistance evolution has been described for various bacterial pathogens, including *E. coli* or *Mycobacterium tuberculosis* (Castro *et al*, 2021; Dubinsky *et al*, 2020; Dupont *et al*, 2017), where infection was suspected to originate from biofilms (Béchon & Ghigo, 2021; Chakraborty *et al*, 2021a; Eldholm *et al*, 2014; Macfarlane & Dillon, 2007; Swidsinski & Lee, 2001). In general, clinical longitudinal studies are often missing, and the link between biofilm infections and within-patients antibiotic resistance could still be underestimated.

Previous *in vitro* studies of the evolution of antibiotic resistance within biofilms used either constant sub-MIC (Ahmed *et al*., 2018; Penesyan *et al*., 2019; Santos-Lopez *et al*., 2019; Trampari *et al*., 2021), step-wise increased concentration starting from sub-MIC (Miller *et al*., 2013; Santos-Lopez *et al*., 2019; Scribner *et al*., 2020) or constant lethal concentration (France *et al*., 2019). Our experimental settings, instead, used periodic treatment with lethal concentrations, above the MPC. In both planktonic and biofilm condition, such periodic regimen generated, at the population level, a low diversity of mutation in genes related to antibiotic recalcitrance, including *fusA* and *sbmA*. These results are in good agreement with the notion that lethal antibiotic concentrations lead to a stringent selection of a few selected mutations increasing the MIC in a single genetic event (Sommer *et al*, 2017). However, the characteristic increased diversity of mutations observed in previous biofilm experimental evolution experiments was, in our case, detected at the allelic level, as in the case of the gene encoding the peptide transporter SbmA, in which up to five mutated alleles could be detected at frequency higher than 5% in the same end-point biofilm population.

Another difference observed between evolved biofilm and planktonic populations was the detection of *fusA* mutations coding for the elongation factor G, an essential protein that facilitates the translocation of the ribosome along the mRNA molecule. *fusA* mutations were found at frequency higher than 5% in 3 out of six biofilm populations exposed to antibiotics. In one of them, a *fusA* mutation was detected early and maintained throughout all evolution cycles. At the clonal level, *fusA* mutations were also detected in planktonic populations, albeit only after concentrating the samples, suggesting that these mutations were also present in planktonic populations but were not selected, while being selected and maintained in biofilm populations. All *fusA* mutations were exclusively located in domains IV and V of the FusA protein. This bias towards these two domains was also observed in other evolution experiments of *E. coli* in the presence of aminoglycosides (Ibacache-Quiroga *et al*., 2018; Jahn *et al*., 2017; Lázár *et al*., 2013). This therefore suggests that the amikacin treatment biased the selection of mutations towards domains IV and V. Consistently, the alignment of all the *E. coli fusA* sequences in the nrprot database of NCBI (2143 sequences) showed that domains I and V are the more variable but that it was not the case of domain IV (supplementary Fig. S11). Therefore, it is very unlikely that unbiased mutations be only found in domains IV and V in our evolved populations (χ2 test, p-value = 3e-06). Interestingly, while we could not precisely correlate these domains IV and V mutations with a precise antibiotic treatment, we identified 11 *fusA* sequences with the A678V mutation and 6 sequences with the P610S mutation (supplementary Table S7) suggesting that the selection of *E. coli fusA* mutations could bear relevance beyond our *in vitro* experiments.

Surprisingly, convergent *fusA* mutations detected in both planktonic and biofilm population of *A. baumannii* and *P. aeruginosa* treated by stepwise increased concentration of tobramycin (Scribner *et al*., 2020) did not display any apparent domain bias contrary to what was observed for *E. coli* in several studies including ours (Ibacache-Quiroga *et al*., 2018; Jahn *et al*., 2017; Mogre *et al*., 2014). The elongating factor G domains IV and V are relatively rigid domains that move together with domain III during the pre- to post-translocation of charged tRNA from the A to the P-site of the ribosome (Lin *et al*, 2015). During the translocation, domain V flips by 180° with a simultaneous 90° self-rotation causing domain IV to project into the decoding center of the 30S ribosome where the anticodon end of the A-site tRNA would be bound. Aminoglycosides bind the A-site decoding center, stabilize the interaction of tRNA in the A-site and ultimately inhibit elongation factor G-catalyzed translocation (Borovinskaya *et al*, 2007; Feldman *et al*, 2010; Tsai *et al*, 2013). One can speculate that *fusA* mutations detected in our study could modify the dynamics of elongation factor G during translocation and/or its affinity with the A-site, thereby modifying binding of aminoglycosides to the decoding center of the 30S ribosome. Alternatively, but not mutually exclusive, such mutations could cause dysregulation of the expression of other factors leading to indirect effect on aminoglycosides activity, as recently shown for a mutation in the *fusA1* gene of *P. aeruginosa*, a hot-spot for mutations in patients with cystic fibrosis (Maunders *et al*, 2020).

One major question raised by our study is why biofilms so rapidly accumulated resistance mutations. One possible explanation obviously lies in the well-described biofilm-associated oxidative and chemical stress response leading to enhanced mutation rate (Boles & Singh, 2008; Driffield *et al*., 2008; Shewaramani *et al*, 2017), also observed in our experimental settings. The high frequency of bacteria possessing different mutations enhancing amikacin’s MIC within biofilm populations could explain their higher capacity to survive lethal amikacin concentration and reflect biofilm propensity to favor clonal interference as compared to planktonic population (Steenackers *et al*., 2016). Biofilm intrinsic tolerance to lethal antibiotic concentration as high as 80xMIC also increases the size of surviving bacteria within biofilms at each cycle of treatment, while planktonic populations were eliminated after three cycles of such a treatment.

This enhancing survival also likely played a role in biofilm capacity to shelter mutants that were early selected for their antibiotic resistance. Interestingly, such relationship between tolerance and resistance, with tolerance promoting the emergence of resistance, has also been demonstrated in planktonic populations treated intermittently with antibiotics (Levin-Reisman *et al*., 2019; Levin-Reisman *et al*., 2017; Windels *et al*., 2019a). However, in our case, tolerance was not provided through selection of *de novo* induced genetic mutations previously reported to increase tolerance in periodically aminoglycoside treated *E. coli* planktonic populations (*i.e.* in *nuoN*, *oppB* or *gadC*) (Van den Bergh *et al*., 2016b), but by the biofilm intrinsic antibiotic tolerance. If these mutations did emerge, the stringency and length of the antibiotic treatment used in our study (24h) could have prevented the selection of such mutations, by contrast with previous studies using shorter treatment (maximum of 8h) (Sulaiman & Lam, 2021).

Studies exploring biofilm evolution under constant antibiotic pressure were rather shown to promote the emergence of mutants with lower resistance level compared to evolved planktonic populations (Ahmed *et al*., 2018; Santos-Lopez *et al*., 2019), which is in stark contrast with the outcomes of our experimental evolution experiments showing that biofilm evolved clones displaying higher resistance than planktonic evolved clones. The use of lethal concentration of antibiotic in our intermittent treatment logically favored the selection for high resistance mutations in both lifestyles (Sommer *et al*., 2017), including *fusA* mutations, conferring resistance to up to 3 times the amikacin MIC. However, these mutations were rapidly counter selected in planktonic populations, while being maintained in biofilms. We showed that *fusA* mutations are associated with a strong fitness cost in planktonic conditions, suggesting that they could be rapidly eliminated in planktonic conditions, notably during the regrowth period without antibiotic. The capacity of structured biofilm environments to maintain high cost mutations was observed in several biofilm evolution studies (Ahmed *et al*., 2020; Ahmed *et al*., 2018; France *et al*., 2019; Santos-Lopez *et al*., 2019), in which rare beneficial mutations could be protected from competitions in biofilm microniches, increasing their odds to be selected and propagated. In support of this hypothesis, it was recently shown that the SOS response induced in *E. coli* biofilm microniches leads to increased rate of mutagenesis (Tlili *et al*, 2021) and that beneficial resistance mutations had reduced probability to be lost by genetic drift in structured environments (Chakraborty *et al*, 2021b). Alternatively, due to diffusion limitation, the concentration of antibiotics could drop more slowly in biofilms than in planktonic conditions. Traces of antibiotics still present in biofilms between each cycle of antibiotics treatment could reduce the fitness cost of resistance mutations by providing a compensatory growth advantage.

Whereas we did not detect mutations increasing antibiotic tolerance, we identified, in addition to *bona fide* aminoglycoside resistance mutations in *fusA* and *sbmA*, several mutations in the *fimH* gene encoding the tip-lectin of the type 1 fimbriae. We recently showed that, in *E. coli* populations grown in dynamic continuous flow conditions, *fimH* is indeed under strong positive selection for mutation promoting non-specific adhesion to surfaces and biofilm formation (Yoshida *et al*., 2022). Mutation in FimH tip-lectin domain identified in the present study not only enhanced *E. coli* biofilm formation, but also its survival to lethal high concentration of amikacin. Such enhanced biofilm capacity has also been observed in *A. baumannii* bead biofilms constantly exposed by sub-MIC or step-wise increased concentration of ciprofloxacin (Santos-Lopez *et al*., 2019), on biofilm formed in silicone tubing treated for 3 days with tetracycline (Penesyan *et al*., 2019), and with *E. faecalis* biofilms exposed to increasing concentration of daptomycin in turbidostats (Miller *et al*., 2013). While our experimental set-up did not expose biofilms to continuous flow, biofilms formed on medical-grade silicone coupons were periodically exposed to washes and flux, probably promoting the selection of biofilm promoting mutations. The selection of mutations promoting biofilm formation could therefore be seen as contributing to a virtuous (or vicious) circle, in which enhanced biofilm formation *de facto* enhances tolerance and survival of biofilm bacteria. This would, in turn, increase the odds for *de novo* genetic resistance mutations to occur.

Together with other works, our study further demonstrates that, depending on the used antibiotic dose or regimen, antibiotic tolerant biofilm environments could promote the worrisome evolution of antibiotic resistance (Ahmed *et al*., 2018; France *et al*., 2019; Miller *et al*., 2013; Penesyan *et al*., 2019; Santos-Lopez *et al*., 2019; Scribner *et al*., 2020; Trampari *et al*., 2021). However, encouragingly, a recent study showed that using short breaks (no more than 4h) between the oxacillin antibiotic treatments of *S. aureus* catheter-associated infection increased the efficacy of the treatment, while preventing the emergence of resistance (Meyer *et al*., 2020). This therefore suggest that improving our current understanding of the evolutionary path taken by biofilms under antibiotic pressure in various regimen and models could lead to new and clinically practical therapeutic options to fight biofilm-related infections.

## MATERIAL AND METHODS

### Bacterial strains and growth conditions

Bacterial strains used in this study are listed in supplementary Table S8. The adherent/invasive AIEC strain *E. coli* LF82 was isolated from a chronic ilea lesion from Crohn’s disease patients (Glasser *et al*, 2001). Overnight cultures were grown at 37°C using Miller’s LB medium (Thermo Scientific, Rochester, NY, USA) under shaking at 180 *rpm* unless otherwise specified. The following antibiotics were used for strain construction: kanamycin (50 µg/mL) and chloramphenicol (25 µg/mL). All compounds were purchased from Sigma-Aldrich (St Louis, MO, USA).

### Strain construction

#### Generation of mutants in *E. coli* LF82

We generated the LF82 mutants used in this study by λ-red linear recombination (Chaveroche *et al*, 2000). When required, antibiotic resistance markers flanked by two FRT sites were removed using the Flp recombinase (Cherepanov & Wackernagel, 1995). We verified the integrity of all cloned fragments, mutations, and plasmids by PCR with specific primers and whole-genome sequencing.

#### Construction of LF82_mars and LF82_GFP

Red (mars) or green fluorescent protein (gfp) encoding genes were inserted on the chromosome of LF82 at the place of the *ampC* gene (supplementary Table S8). In brief, plasmid pZE1RGFP-KmFRT was constructed by amplification of KmFRT region from plasmid pKD4 and cloning between SalI-HindIII sites of plasmid pZE1RGFP. Plasmid pZE1R-mars-KmFRT was constructed by cloning the pZS*2R-mars plasmid’s KpnI-BamHI digested region containing the mars open reading frame between the KpnI-BamHI sites of pZE1RGFP-KmFRT plasmid. Then the DNA fragments of mars-KmFRT and gfp-KmFRTwere amplified from, respectively, pZE1R-mars-KmFRT and pZE1RGFP-KmFRT using long recombining primers (supplementary Table S9). The resulting PCR products, which have approximately 50 bp-long regions of homology upstream of fluorescent genes and downstream of KmFRT, were introduced at the *ampC* gene by λ-red recombination into strain LF82 using pKOBEG plasmid (Chaveroche *et al*., 2000), respectively. Then, Δ*ampC::mars-*KmFRT and Δ*ampC::GFP-*KmFRT mutations were introduced into strain LF82 by P1vir phage transduction, in which the Flp recombinase was used to remove the kanamycin marker using pCP20 plasmid (Cherepanov & Wackernagel, 1995).

#### Deconstruction of the fusA G604V allele in clone 219

The CmFRT cassette (*cat* gene) was amplified from strain MG1655*ΔyfcV-P::*CmFRT (Korea *et al*, 2010) using long recombining primers (supplementary Table S8, S9). The resulting PCR product, which has approximately 50 bp-long regions of homology upstream and downstream of CmFRT start and stop codons, was introduced just after the *tufA* gene that follows the *fusA* gene by λ-red recombination into strain LF82_*mars* using pKOBEGA plasmid (Chaveroche *et al*., 2000). This generates the strain LF82_mars, *fusAwt-*CmFRT. A DNA fragment encompassing *fusA_wt-*CmFRT was amplified from strain LF82_*mars*, *fusAwt*-CmFRT using the primers, Up.tufA-CmFRT-5 and Down.tufA-CmFRT-3 (supplementary Table S9). The *fusA* G604V allele of the clone 219 corresponding to LF82_*mars, fusA_G604V, fimH_L79R, LF82_RS22000_I255I* was deconstructed back to wt by it by the *fusA_wt*-CmFRT region using λ-red recombination with the pKOBEGA plasmid.

### Amikacin concentration dependent killing of *E. coli* LF82 in biofilm and planktonic conditions

LF82 biofilm- and planktonic- cells were treated with or without AMK at 1- to 80-fold MICs (1, 5, 10, 20, 50, and 80-fold MICs) for 24 h, respectively. The survival rate of bacteria under biofilm or planktonic conditions after AMK treatment was calculated as the ratio of the number of survived cells after the treatment per the total number of bacterial cells before the treatment by performing CFU counts using bacterial suspensions.

#### i) Biofilm cells on silicone coupons

LF82 biofilms were grown on 30 silicone coupons sheeted in a well of a 6-well plate (Techno Plastic Products AG) and treated with AMK. Briefly, silicone sheets (Silicone elastomer membrane 7-4107, Dow Corning Corporation, Midland, MI, USA) were cut out into round pieces at a 5 mm diameter using a biopsy punch (Kai Medical, Seki City, Japan). Then, cut coupons were placed on a well in 6-well plate (30 coupons per well), and the plate was sterilized with ethylene oxide. LF82 overnight culture was diluted to an optical density at 600 nm (OD_600_) of 0.05 in LB medium, and 5 mL of the diluted culture was poured into the well. Biofilms were grown at 37 °C for 24 h under static condition. After 24-h incubation, the exhausted medium containing floating cells was removed from the well using an aspirator (VACUSIP; Integra Biosciences, Hudson, NH, USA), and the well was gently washed once with 5 mL of phosphate buffered saline (PBS, Lonza, Rockland, ME, USA) to remove un-adherent floating bacteria. Six coupons were aseptically collected using sterilized forceps and each dipped in 500 μL of LB medium in a microtube as before treatment coupons. The remaining biofilm cells in the well were treated with AMK dissolved in 5 mL of fresh LB medium at 37°C for 24 h. After removing the supernatant, the well was rewashed, and six coupons were collected each in 500 μL of LB medium as treatment coupons. Each microtube containing one coupon (before treatment coupons and treatment coupons) was vigorously vortexed for 1 min and sonicated for 10 min using an ultrasonic bath (Branson 5800, Branson Ultrasonics, CT, USA) to detach biofilm cells from coupons and prepare bacterial suspension. The bacterial number of the suspension was confirmed by performing CFU counts.

#### ii) Planktonic cells

LF82 was incubated in a test tube containing 3 mL of fresh LB medium at 37°C for 24 h under shaking condition. After 24 h incubation, LF82 culture was treated with AMK at 37 °C for 24 h under shaking condition by directly adding several concentrations of AMK solution in the test tube. Before and after the AMK treatment, 200 μL of the culture was collected in a microtube. Then, bacterial pellets were harvested by centrifuging at 15,000 *rpm* for 3 min at 25 °C, washed once with 200 μL of PBS, and suspended in 200 μL of LB medium to prepare bacterial suspension. The bacterial number of the suspension was confirmed by performing CFU counts. The test was repeated six times for each concentration.

### Mutation Prevention Concentration determination

We determined the MPC value that prevents the growth of at least 1.0 × 10^10^ bacteria of LF82, according to previous literature (Drlica, 2003). Briefly, 100 mL of LF82 overnight culture was centrifuged at 4,500 *rpm* for 10 min at 25°C. After removing the supernatant, bacterial pellets were suspended in 10 mL of fresh LB and concentrated to approximately 2.0 × 10^10^ bacteria/mL. The bacterial number of the plated suspension was confirmed by performing CFU counts on an LB agar plate after 24-h incubation. Subsequently, 100 μL of the bacterial suspension were plated on LB agar plates with or without AMK at 1- to 6-fold MIC and incubated at 37°C for 72 h. Ten replications were made for each concentration of AMK to evaluate the growth for a total of approximately 2.0 × 10^10^ bacteria. The MPC corresponds to the AMK concentration of the plate where no growth was observed.

### Experimental evolution in biofilm and planktonic populations under intermittent amikacin exposure with lethal concentrations

Evolution experiments, in which biofilm and planktonic populations were intermittently treated with or without AMK at 5- or 80-fold MICs, were performed for 10 cycles. In both conditions, one cycle consists of a 24 h AMK treatment and two 24 h regrowth steps in absence of AMK, with washing steps between 24 h treatment/incubation steps (Steps 2–6 in Fig. 1 were defined as a series of one cycle). Evolved populations and end-point clones obtained from experiments were stocked as glycerol stock at -80°C and characterized later by MIC testing, the frequency of AMK resistant mutants, survival rate against AMK treatment, and whole-genome sequencing, if necessary.

#### i) Biofilm condition

First, LF82_*GFP* and LF82_*mars* from glycerol stocks were grown overnight in LB medium under shaking condition and were adjusted to OD_600_ of 0.05 in LB medium. After mixing the adjusted cultures of LF82_*GFP* and LF82_*mars* at a 1:1 ratio, 5 mL of the mixture was poured into a total of 9 wells (3 wells of a 6-well plate for each of the following conditions: control, 5-, or 80-fold MIC treatments). Each well was sheeted with 30 silicone coupons and incubated under static condition at 37°C for 24 h to form biofilm on silicone coupons. Then, the exhausted medium containing floating cells was removed from the well, and the well was gently washed twice with 5 mL of LB medium as described above (hereafter, called washing step). Biofilms were incubated with 5 mL of fresh LB medium for an additional 24 h (in total of 48 h, Fig. 1A, step 1). After performing washing steps twice, one coupon was collected from each of the 9 wells in 500 μL of LB medium into a microtube (Fig. 1A step 2; hereafter, called coupon sampling). Next, biofilm cells contained in 3 wells were incubated with LB (control), with 5- or 80-fold MICs of AMK dissolved in 5 mL of LB medium (Fig. 1A step 3) under static condition at 37°C for 24 h. After removing the supernatant containing AMK and performing washing steps twice (Fig. 1A step 4), one coupon was collected in each well (Fig. S1A step 5). Then, 5 mL of LB medium was poured into the well to allow survived biofilm cells to regrow. Next, the survived biofilm cells were incubated for 24 h at 37°C, and coupon sampling was performed in each well after removing the supernatant and performing washing steps twice (Fig. 1A step 6). Finally, the biofilm cells were incubated with 5 mL of LB medium for an extra 24 h (in total of 48 h) before the subsequent AMK exposure (Fig. 1A step 2). Steps 2-6 were defined as a series of cycles, and the experiment is performed for 10 cycles. In each sampling step, bacterial suspensions were prepared by detaching biofilm cell from coupons as mentioned above. Then, the bacterial number of the suspension was confirmed by performing CFU counts and stocked as glycerol stock at -80°C and characterized later. From CFU counts of last cycle at step 5, some colonies were randomly picked as endpoint clones and used for characterization.

#### ii) Planktonic condition

First, LF82_*GFP* and LF82_*mars* from glycerol stocks were grown for 24 h in LB medium at 37°C for 24 h under shaking condition. The two cultures were adjusted to OD_600_ of 0.01 in LB medium into test tubes and incubated for an extra 24 h (in total of 48h, Fig. 1B step 1). LF82_*GFP* and LF82_*mars* cultures were mixed at a 1:1 ratio, after diluting each culture to OD_600_ of 2.0. Then, 200 μL of the mixed culture was collected and stored for further analyses (Fig. 1B step 2). Next, 3 mL of the mixed culture was transferred into 9 test tubes and 3 cultures were incubated without AMK, 3 with AMK at 5-MICs and the remaining 3 cultures with AMK at 80-fold MICs (Fig. 1B step 3). After 24 h of incubation under shaking condition, 1 mL aliquots of each of the 9 cultures were centrifuged, and bacterial pellets were harvested, washed twice, and suspended in 1 mL of LB medium (Fig. 1B step 4). Then, 500 µL aliquots of the 9 suspensions were stored for further analyses (hereafter, called sampling step, Fig. 1B step 5). Next, 30 μL of each suspension was inoculated in 3 mL of fresh LB medium (*i.e.* 100-fold dilution) and incubated for 24 h at 37°C under shaking condition. Subsequently, after performing bacterial collection washing- and sampling-steps (Fig. 1B step 6), 1/100-fold suspension was inoculated in 3 mL of LB medium and incubated before the subsequent AMK treatment (Fig. 1B step 2). Steps 2-6 were defined as a series of cycles, and the experiment is performed for 10 cycles. The last cycle was stopped at step 5. In each collected culture, the bacterial number was confirmed by performing CFU counts and stocked as glycerol stock at - 80°C and characterized later. From CFU counts of last cycle at step 5, some colonies were randomly picked as endpoint clones and used for characterization.

### Evaluation of the numbers of generation propagated during experimental evolution experiments

The numbers of generation during which bacteria were propagated were evaluated using CFU counting performed at step 2, 5 and 6 of each cycle of evolution, both for biofilm and planktonic populations (Fig. 1). For biofilms, generation time was calculated by calculating n1= LOG(CFUstep6;2)-LOG(CFUstep5;2) and n2= LOG(CFUstep2;2)-LOG(CFUstep6;2). For planktonic populations, because of two steps where 1/100-fold dilution was performed, generation time was calculated by calculating n1= LOG(CFUstep6;2)-LOG(CFUstep5/100;2) and n2= LOG(CFUstep2;2)-LOG(CFUstep6/100;2). Therefore, for each cycle the total number of generation times was n1+n2. Calculated generation times were then cumulated over the cycles (supplementary Table S1).

### MIC determination

We determined the MIC values of amikacin (AMK) (Sigma-Aldrich) or rifampicin (RIF) (Sigma-Aldrich) for each strain using the agar dilution method (Clinical_and_Laboratory_Standard_Institute, 2022) with some minor modifications. Briefly, strains were inoculated into test tubes containing LB medium and incubated overnight at 37°C. After diluting the overnight culture to approximately 1.5 × 10^8^ CFU/mL into fresh LB medium, 0.5 μL of the diluted culture was inoculated to LB agar plates containing AMK or RIF at different concentrations using a Microplanter inoculator (Sakuma Factory, Tokyo, Japan). The concentration of AMK was adjusted at a finer level than 2-fold increment (specifically, 4, 8, 12, 16, 20, 24, 32, 48, 64, 128, 256 µg/mL) to increase the sensitivity for finding the susceptibility changing. The concentration of RIF was set conducted in 2-fold increments according to the normal agar dilution method.

### Frequency of AMK-resistant mutants

Aliquots of frozen stocks of evolved biofilm- and planktonic-populations obtained from step 5 (Fig. 1A & B) in each cycle were directly inoculated to LB medium into test tubes and incubated overnight under shaking condition at 37°C. As described above, CFU counts for these cultures were performed on LB agar plates with or without AMK at 1-, 2- or 4-fold MICs. The frequency of AMK resistant-mutants was calculated for each AMK concentration as follows:

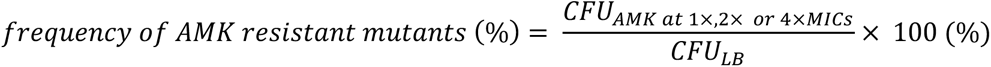

where *CFU*_*AMK* at 1×,2×,or 4×*MIC*_ is the CFU number of each population grown on LB agar plates containing AMK at 1-, 2- or 4-fold MICs, and *CFU_LB_* is the one grown on LB agar plate.

### Survival of endpoint clones

#### i) Biofilm cells on silicone coupons

Percentages of survival of endpoint clones from biofilm evolution on biofilms against 5- and 80-fold MIC of amikacin were evaluated as described above. Briefly, the randomly picked clones were overnight cultured and diluted to an optical density at 600 nm (OD_600_) of 0.05 in LB medium, and 5 mL of the diluted culture was poured into the well with silicone coupons. Biofilms were grown at 37 °C for 24 h under static condition. After 24-h incubation, the exhausted medium containing floating cells was removed from the well using an aspirator (VACUSIP; Integra Biosciences), and the well was gently washed once with 5 mL of PBS (Lonza, Rockland) to remove un-adherent floating bacteria. Three coupons were aseptically collected using sterilized forceps and each dipped in 500 μL of LB medium in a microtube as before treatment coupons. The remaining biofilm cells in the well were treated with AMK dissolved in 5 mL of fresh LB medium at 37°C for 24 h. After removing the supernatant, the well was rewashed, and three coupons were collected each in 500 μL of LB medium as treatment coupons. Each microtube containing one coupon (before treatment coupons and treatment coupons) was vigorously vortexed for 1 min and sonicated for 10 min using an ultrasonic bath (Branson 5800, Branson Ultrasonics, CT, USA) to detach biofilm cells from coupons and prepare bacterial suspension. The bacterial number of the suspension was confirmed by performing CFU counts.

#### ii) Planktonic cells

Percentages of survival of endpoint clones from biofilm and planktonic cell evolution of planktonic cells against 5- and 80-fold MIC of amikacin were evaluated as described above. Briefly, strains were incubated in a test tube containing 3 mL of fresh LB medium at 37°C for 24 h under shaking condition. After 24 h incubation, LF82 culture was treated with AMK at 37 °C for 24 h under shaking condition by directly adding 5- or 80-fold MIC of AMK in the test tube. Before and after the AMK treatment, 200 μL of the culture was collected in a microtube. Then, bacterial pellets were harvested by centrifuging at 15,000 *rpm* for 3 min at 25 °C, washed once with 200 μL of PBS, and suspended in 200 μL of LB medium to prepare bacterial suspension. The bacterial number in the suspension was confirmed by performing CFU counts. The test was repeated six times for each concentration.

### Genome sequencing and analysis

The genomic DNAs were extracted from LF82 mutants (generated constructs and end-point clones of evolution experiments) and evolved populations obtained from experimental evolution. The glycerol stock was scraped, incubated in 1 mL of LB and incubated for 16h at 37°C under shaking conditions. The cells were then pelleted and used for genome extraction using the Wizard Genomic DNA Purification Kit (Promega, Madison, WI, USA) and Qiaquick PCR purification kit (Qiagen, Hilden, Germany). The extracted genomic DNAs were prepared for whole genome sequencing using the Nextera XT DNA library preparation kit (Illumina, San Diego, CA, USA) and sequenced on Hiseq and Miseq platform (Illumina) with an average depth of 150x. Trimmed sequence reads were analysed by BreSeq version 0.35.0 (Deatherage & Barrick, 2014) to detect genetic variants using default parameters for clones and by adding the -p flag for the populations. No chromosomal rearrangements or movement of insertion elements were detected as selected by the applied treatment. The Muller Plots were inferred and produced using the lollipop tool (https://github.com/cdeitrick/Lolipop) as well as the ggmuller R package 0.5.3 (Robert Noble (2019). ggmuller: Create Muller Plots of Evolutionary Dynamics; https://CRAN.R-project.org/package=ggmuller).

### Bioinformatic analysis of FusA sequences

All *E. coli* FusA proteins contained in the nrprot database of NCBI (version 2019-04-09) were identified by BLASTp (blast+ version 2.2.31, (Camacho *et al*, 2009)) and only hits with an e-value lower than 1e^-10^ and sequence identity higher than 70% were kept. The resulting 2143 sequences were aligned using mafft version 7.407 (Yamada *et al*, 2016) with G-INS-I option and the alignment was used to compute entropy in each structural domain of the protein using the HIV sequence database website tool (https://www.hiv.lanl.gov/content/sequence/ENTROPY/entropy.html).

The same alignment was also screened to identify mutations relatively to our reference sequence from LF82 as in (Yoshida *et al*., 2022).

### Growth capacity

An overnight culture of each strain was diluted to OD_600_ of 0.05 in fresh LB medium. Two hundred-μL aliquots were inoculated in a 96-well plate. The plates were then incubated in a TECAN Infinite M200 Pro spectrophotometer (Männedorf, Switzerland) for 20 hours at 37°C with shaking of 2 mm amplitude. The absorbance of each culture at 600 nm was measured every 15 min. Growth curve of each strain was measured using 3 biological replicates.

The R package GrowthCurver (Sprouffske & Wagner, 2016) was used to infer the growing capacities for all strains based on the growth curves.

### Competition assay for *fusA* and *sbmA* mutants in the presence or absence of AMK

Competition assay for *fusA* and *sbmA* mutants were performed according to a previously published method (Van den Bergh *et al*., 2016b), with some modifications. Briefly, LF82_*mars*, LF82_*GFP*, and evolved strains (*fusA* and *sbmA* mutants) tagged with *mars* were grown overnight in LB at 37°C. All overnight cultures were diluted to OD_600_ of 2.0. Then, LF82_*mars* and mars-tagged evolved strains were mixed with LF82_*GFP* at a 1:1 ratio. The mixture of LF82_*mars* and LF82_*GFP* was used as a reference competition. Before starting competition assays, each mixture was verified to be in 1:1 ratio by MACSQuant VYB flow cytometer (Miltenyi Biotec, Bergisch Gladbach, Germany). The experiments i) and ii) were performed using 3 biological replicates in each competition.

The relative fitness of each evolved strain was calculated as follows:

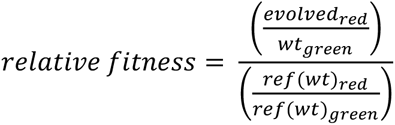

where *wt_green_* and *evolved_red_* are the cell proportions of the LF82_*GFP* and mars-tagged evolved strains in the competition mixture, and *ref(wt)_green_* and *ref(wt)_red_* are the ones of LF82_*GFP* and LF82_*mars* in the reference competition mixture, respectively.

#### i) Competition assay without AMK treatment

Each OD_600_ 2 adjusted mixture was diluted to OD_600_ = 0.02 in 3 mL of LB medium and incubated 24h at 37°C under shaking condition at 180 *rpm*. The relative amount of green and red cells was analysed using flow cytometer to calculate the fitness cost for each evolved strain in the absence of AMK treatment.

#### ii) Competition assay with AMK treatment

Three mL of each mixture at OD_600_ of 2.0 was treated with 5-fold MIC AMK for 24 h at 37°C under shaking condition at 180 *rpm*. Then, 1 mL of each mixed culture was collected in a microtube. Next, bacterial pellets were harvested by centrifuging at 15,000 *rpm* for 3 min at 25 °C, washed twice with 1 mL of LB, and suspended in 1 mL of fresh LB medium to prepare bacterial suspension. Next, 30 μL of the suspension was inoculated in 3 mL of fresh LB medium and incubated 24h at 37°C under shaking condition. Finally, the relative amount of green and red cells was analysed using flow cytometer to calculate the fitness cost for each evolved strain in the presence of AMK treatment.

### Bacterial number and biofilm amount of *fimH* mutants formed on silicone coupons

Overnight cultures of LF82_*mars* and *fimH* mutants grown in LB at 37°C were diluted to OD_600_ of 0.05. Five mL of the diluted culture was inoculated in a well containing 24 silicone coupons and incubated for 3 h at 37 °C under static condition. Then, the supernatant was removed from the well, and the well was gently washed with 5 mL of LB medium once to remove un-adherent bacteria. Next, 5 mL of fresh LB medium was newly added to the well and incubated for 21 h. Then, the exhausted medium containing floating cells was removed, and the well was washed with 5 mL of LB medium twice. Experiments *i)* and *ii)* were performed using 3 biological replicates in each strain.

#### i) CFU counts of biofilm cells on silicone coupons

A total of 6 coupons were randomly collected in 3 microtubes containing 500 μL of LB medium (*i.e.* 2 coupons/microtube). Bacterial suspensions were prepared to perform CFU counts as described above.

#### ii) Cristal violet assay to quantify biofilm amount on silicone coupons

After collecting the 6 coupons for CFU counts, 6 mL of 1% crystal violet (CV) solution (Sigma-Aldrich) was added to the well and incubated for 15 min at 25 °C. The well stained with CV was gently washed with 7 mL of PBS three times to remove excess CV, and the plate was dried up for 1 day in a chemical hood. On the next day, a total of 18 coupons were randomly collected in 6 microtubes (*i.e.* 3 coupons/microtube), which contains 810 μL of the mixed solution of ethanol/acetone at 80%:20% ratio. Coupons were suspended in the solution for 15 min to allow the CV stain to be dissolved. After transferring 100 μL of dissolved CV solution in a 96-well plate, the OD_570_ value was measured to quantify the CV solution using a multimode plate reader (Tecan Infinite M200 PRO). As the background control, 6 coupons sheeted in a well containing 5 mL of LB medium without bacteria were also treated similarly.

### Survival rate of biofilm cells of *fimH* mutants formed on silicone coupons after AMK treatment

Similar to the above, 5 mL of the diluted culture at OD_600_ of 0.05 was inoculated in the well containing 8 coupons and incubated for 3 h at 37 °C. After washing the well once, 5 mL of fresh LB medium was newly added in the well and incubated for 21 h at 37°C. Then, after washing the well twice, 4 coupons were collected in one microtube containing 500 μL of LB medium (*i.e.* 4 coupons/microtube). Then, 5 mL of LB medium containing AMK at 80-fold MIC was added to the well and incubated for 24 h at 37°C. Next, after washing the well twice to remove AMK, the remaining 4 coupons were collected in one microtube containing 500 μL of LB medium (*i.e.* 4 coupons/microtube). Bacterial suspensions were prepared from microtubes containing coupons, as mentioned above. Subsequently, CFUs were quantified on LB agar plates in the same way as described above. Finally, the survival rate of each strain was calculated as follows:

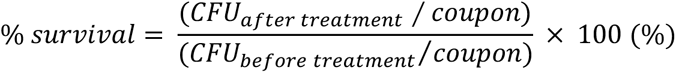

where *CFU_after treatment_* is the bacterial number of the suspension for 4 coupons collected from the well after AMK treatment, and *CFU_before treatment_* is the one for 4 coupons collected from the same well before AMK treatment, respectively. This experiment was performed using 9 biological replicates for each strain.

### Frequency of rifampicin resistant mutants in biofilm and planktonic cells

As mentioned above, after forming 24 h LF82 biofilms on silicone coupons in LB medium in a well of a 6-well plate, a bacterial suspension of biofilm cells was prepared. For planktonic cells, after growing LF82 in a test tube in LB at 37°C for 24 h, 2 mL aliquots of planktonic culture were harvested, washed, and suspended in 120 uL LB medium. Then, the bacterial numbers of each suspension from biofilm and planktonic conditions were confirmed by CFU counts on LB agar plates. Additionally, 100 μL of each bacterial suspension was plated on LB agar plates with RIF at 4-fold MIC (MIC of RIF against LF82 is 16 μg/mL). After 24 h incubation, CFUs were quantified from each plate. The frequency of RIF-mutants was calculated as the ratio of the bacterial number on LB agar plate with RIF at 4-fold MIC per that the one on LB agar plate without RIF.

## DATA AVAILABILITY

All sequencing reads were deposited in NCBI under the BioProject accession number PRJNA833264. The code used in this study are available at https://github.com/Sthiriet-rupert/Usui_Ecoli_Amk.

## ACKNOWLEDGEMENTS

We thank Dr. Olaya Rendueles and Dr David Lebeaux for critical reading of the manuscript. We are grateful to Dr. Olaya Rendueles for the initial help with the analysis of the mutations. This work was supported by the French National Research Agency (ANR), project EvolTolAB (ANR-18-CE13-0010), by the French government’s Investissement d’Avenir Program, Laboratoire d’Excellence “Integrative Biology of Emerging Infectious Diseases” (grant n°ANR-10-LABX-62-IBEID) and by the Fondation pour la Recherche Médicale (grant DEQ20180339185). S.T.-R was supported by the French National Research Agency (ANR), project EvolTolAB (ANR-18-CE13-0010).

## AUTHOR CONTRIBUTIONS

C.B. and M.U. designed the experiments. M.U., Y.Y. and S.T.-R. performed the experiments. C.B., M.U., Y.Y, S.T.-R. and J.-M.G. analysed data. C.B. wrote the manuscript with significant help of M.U., Y.Y, S.T.-R. and J.-M.G.

