## Supplementary material for "Intermittent antibiotic treatment of bacterial biofilms favors the rapid evolution of resistance": Usui_et_al-2022-SuppFig.pdf

**SUPPLEMENTARY FIGURES**

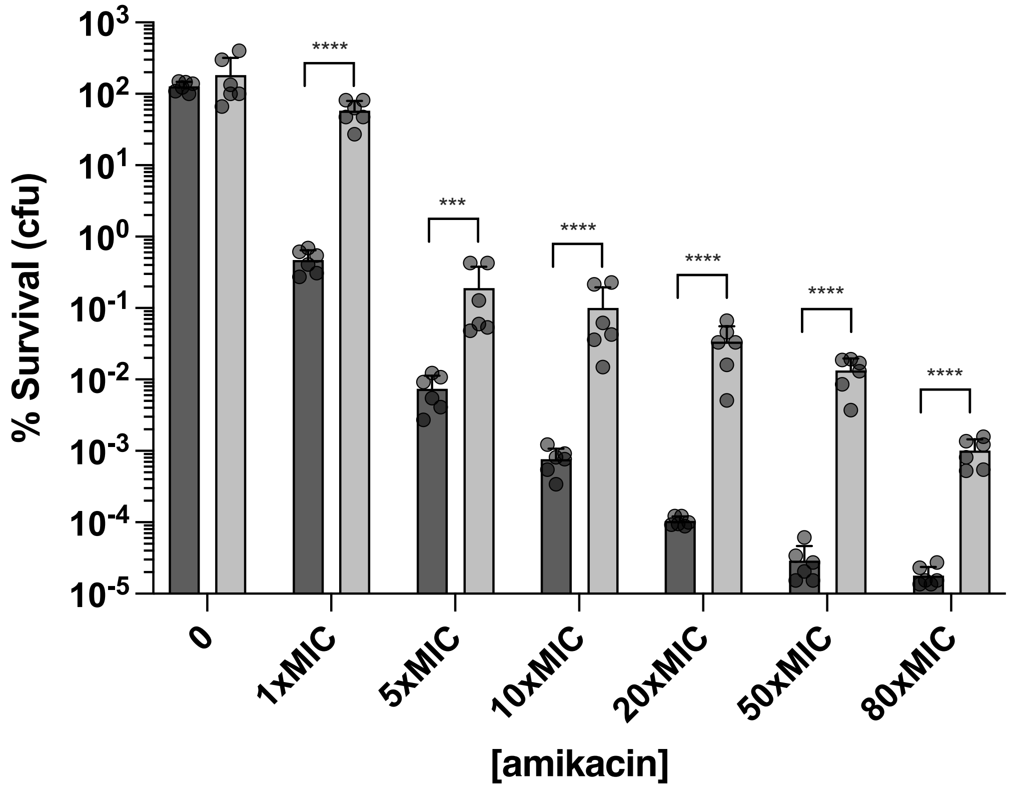


**Supplementary Fig. S1. Amikacin concentration dependent killing of *E. coli* LF82 in planktonic and biofilm conditions.** LF82 biofilms were grown on silicone coupons during 24h at 37°C under static conditions. LF82 planktonic bacteria were grown during 24h at 37°C under agitation. Biofilms (light grey) and planktonic cells (dark grey) were then exposed to increasing concentration of amikacin for 24h, after which silicone coupons or aliquots of planktonic cultures were collected, washed, serially diluted and plated for CFU counting and estimation of the survival rate. Each bar represents the mean of six biological replicates with standard error of the mean. Data were log10 transformed before being submitted to unpaired two-tailed t-test with Welch's correction. *P ≤ 0.05; **P ≤ 0.01; ***P ≤ 0.001, ****P ≤ 0.0001.

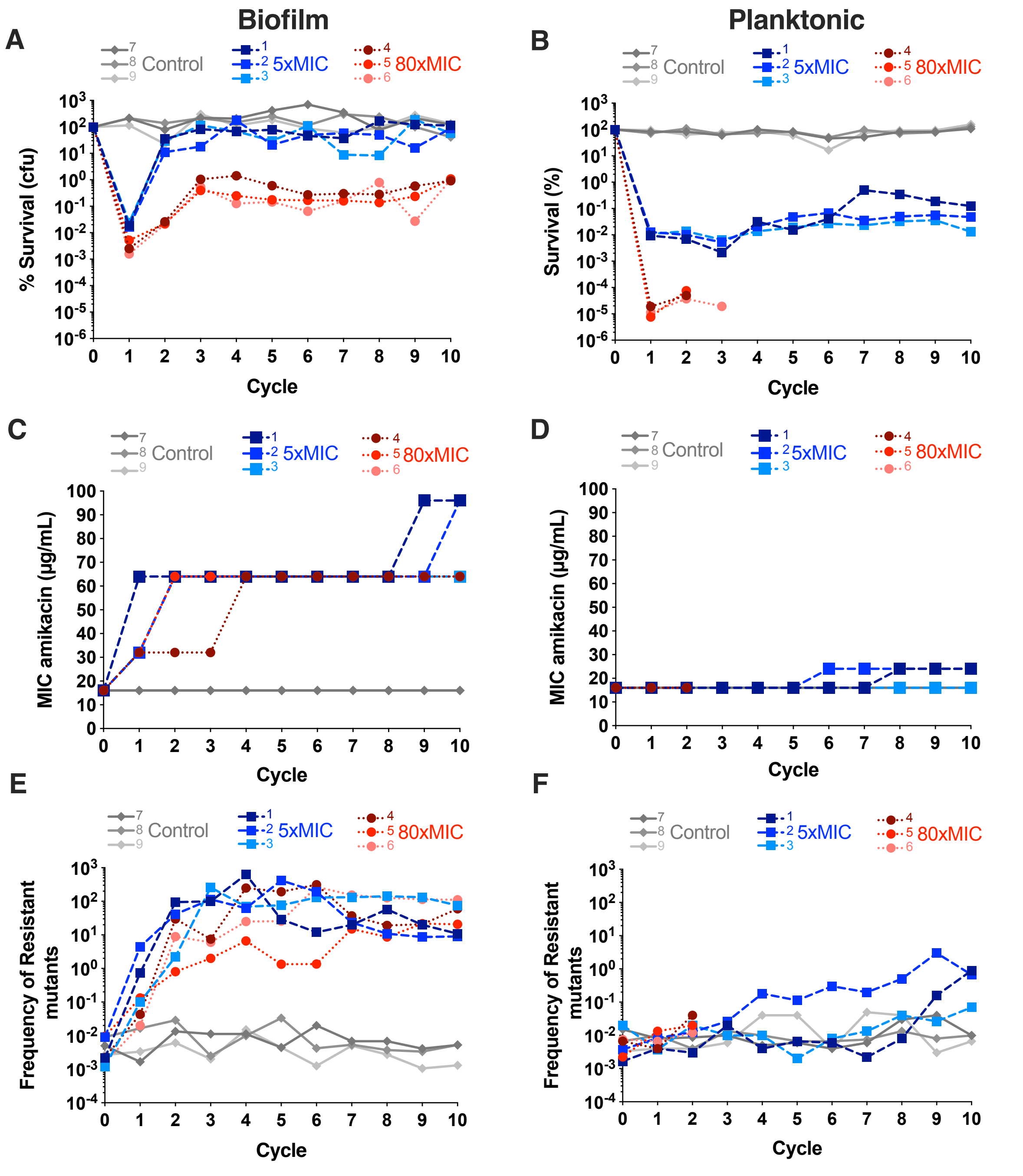


**Supplementary Fig. S2. Evolution of *E. coli* under lethal antibiotic 24h intermittent treatment of amikacin as in Fig. 2 with independent evolution replicates plotted.** Evolution in (A, C, E) Biofilms and (B, D, F) Planktonic**.** In (A, B) are represented the percentage of survival after each cycle of evolution at step 5 for biofilms and planktonic as compared to population before each cycle of treatment at step 2. Percentages of survival were determined used CFU counting. The three independent evolution were performed in parallel for each condition (no treatment= control in grey, 5x amikacin MIC in blue, 80x amikacin MIC in red). In (C, D), each population sampled at the end of each cycle was evaluated for its MIC to amikacin using the agar dilution method. In (E, F), each population sampled at the end of each cycle was plated on LB plate with or without 1x amikacin MIC (or 2xMIC and 4xMIC see supplementary Fig. S4). Frequency of resistant mutants was calculated as the $CFU_{1xMIC}/CFU_{LB}$. In (E, F), the representation is different from the one of Fig. 2 to facilitate reading.

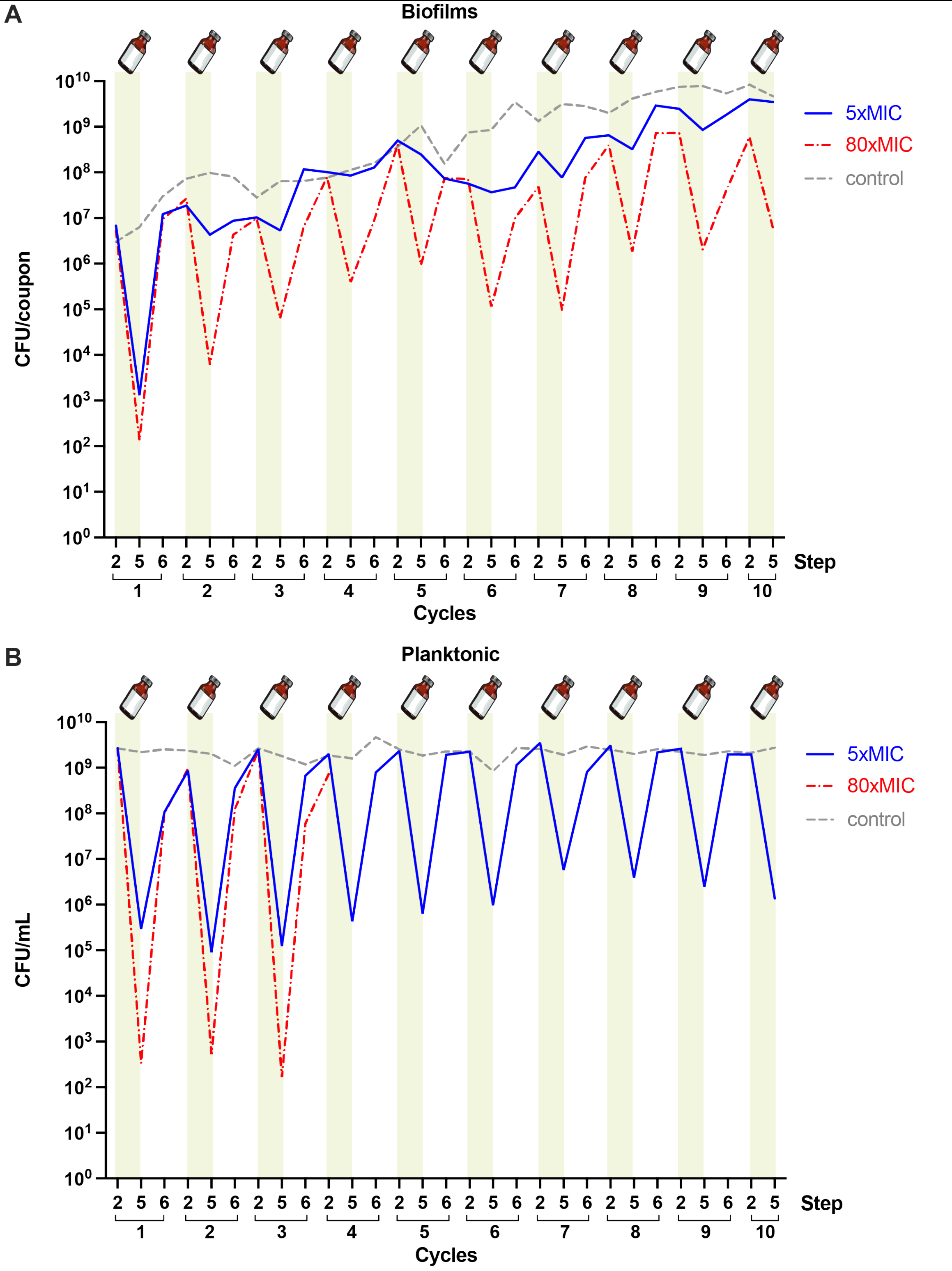


**Supplementary Fig. S3. Evolution of colony forming units at each sampling step over the 10 cycles of the experiments**, in Biofilms (A) and in Planktonic (B)**.** CFU counting correspond to sampling for both biofilm and planktonic at steps 2, 5 and 6 for cycles 1−9 then steps 2 and 5 for cycle 10 (see Fig. 1). Three independent evolutions were performed in parallel for each condition (no treatment = control, 5x amikacin MIC, 80x amikacin MIC). At each step we represented the means of the CFU counts per coupon (biofilm) or mL (planktonic) of the three experiments for control, 5x amikacin MIC, 80x amikacin.

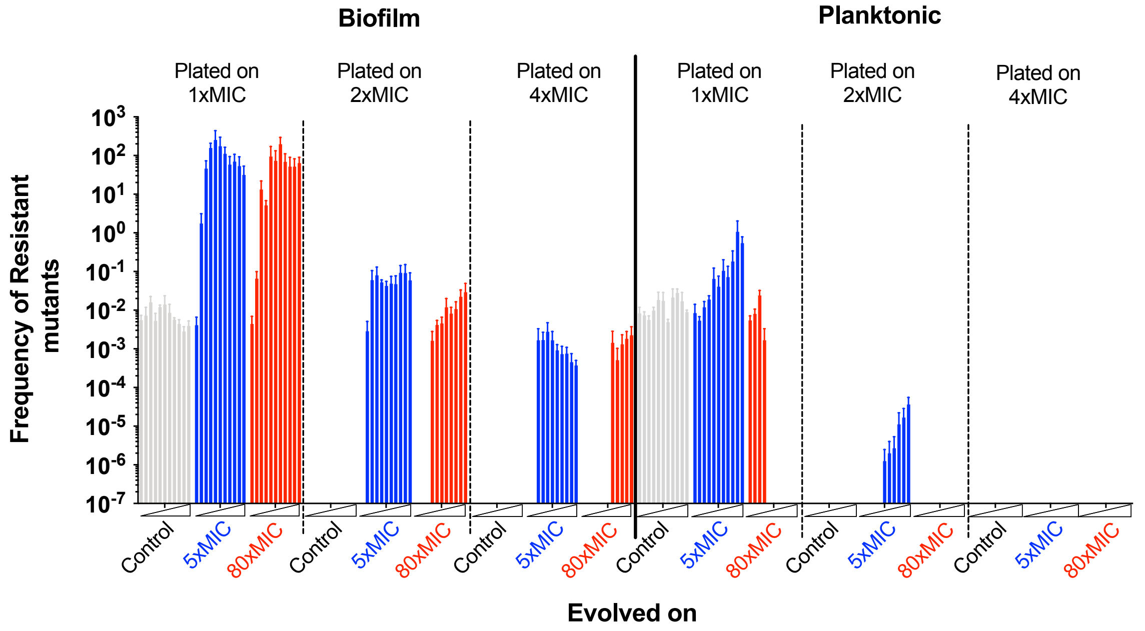


**Supplementary Fig. S4. Frequency of resistant mutants growing on 1xMIC, 2xMIC or 4xMIC of amikacin over cycles of 24h intermittent treatment.** Each 24h intermittently treated biofilm and planktonic population sampled at the end of each cycle was plated on LB plate with or without 1xMIC, 2xMIC or 4xMIC (1xMIC results are duplicated from Fig. 2E and F and presented here for comparison with 2xMIC and 4xMIC). Frequency of resistant mutants was calculated as the $CFU_{1xMIC\ or\ 2xMIC\ or\ 4xMIC}/CFU_{LB}$.

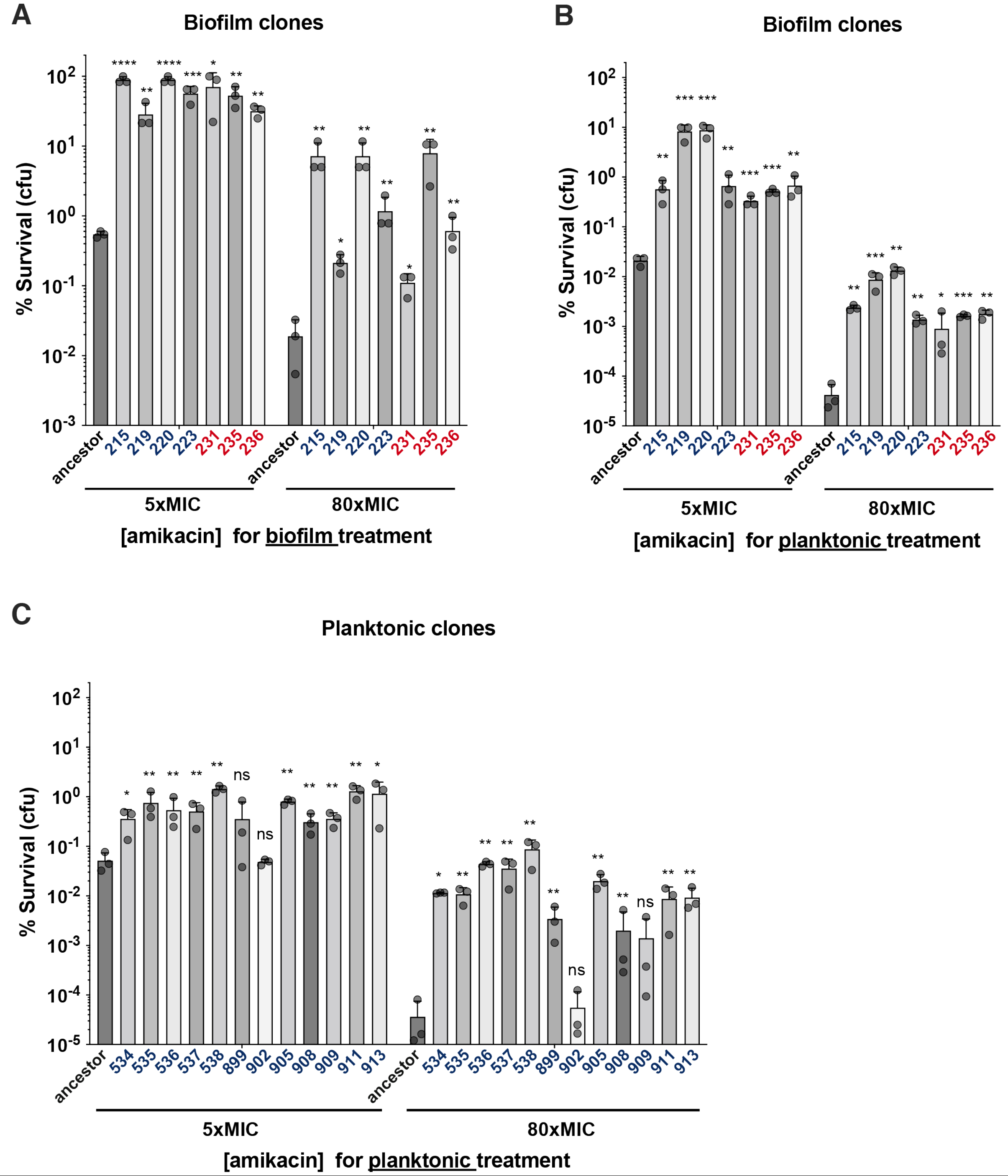


**Supplementary Fig. S5. Survival of endpoint clones from biofilm and planktonic evolved populations.** (A) % of survival of biofilm evolved cycle 10 clones when grown as biofilm on silicone coupons and treated by amikacin. Cycle 10 clones evolved in biofilms in presence of 5x amikacin MIC (in blue) or 80x amikacin MIC (in red) were grown as biofilm during 24h and treated or not for 24h by either of 5x amikacin MIC or 80x amikacin MIC and alive bacteria were quantified by counting colony forming units for each coupon. Survival was expressed as % of survival of the treated samples (n=3 for each clone) as compared to the non-treated samples (n=3 for each clone). (B) % of survival of biofilm evolved cycle 10 clones when grown as planktonic cultures and treated by amikacin. Cycle 10 clones evolved in biofilms in presence of 5x amikacin MIC (in blue) or 80x amikacin MIC were grown as stationary phase cultures and treated or not for 24h by either 5x amikacin MIC or 80x amikacin and alive bacteria were quantified by counting colony forming units. Survival was expressed as % of survival of the treated samples (n=3 for each clone) as compared to the non-treated samples (n=3 for each clone). (C**)** % of survival of planktonic evolved cycle 10 clones when grown as planktonic cultures and treated by amikacin. Cycle 10 clones evolved in planktonic in presence of 5xMIC (in blue) of amikacin were grown as stationary phase cultures and treated or not for 24h by either 5xMIC or 80xMIC of amikacin and alive bacteria were quantified by counting colony forming units. Survival was expressed as % of survival of the treated samples (n=3 for each clone) as compared to the non-treated samples (n=3 for each clone). Data were log10 transformed before being submitted to multiple unpaired t tests with Welch correction and Holm-Šídák method for multiple analysis correction. *P ≤ 0.05; **P ≤ 0.01; ***P ≤ 0.001, ****P ≤ 0.0001 and ns non-significant. These different clones have been sequenced and their associated mutations are described in supplementary Table S4.

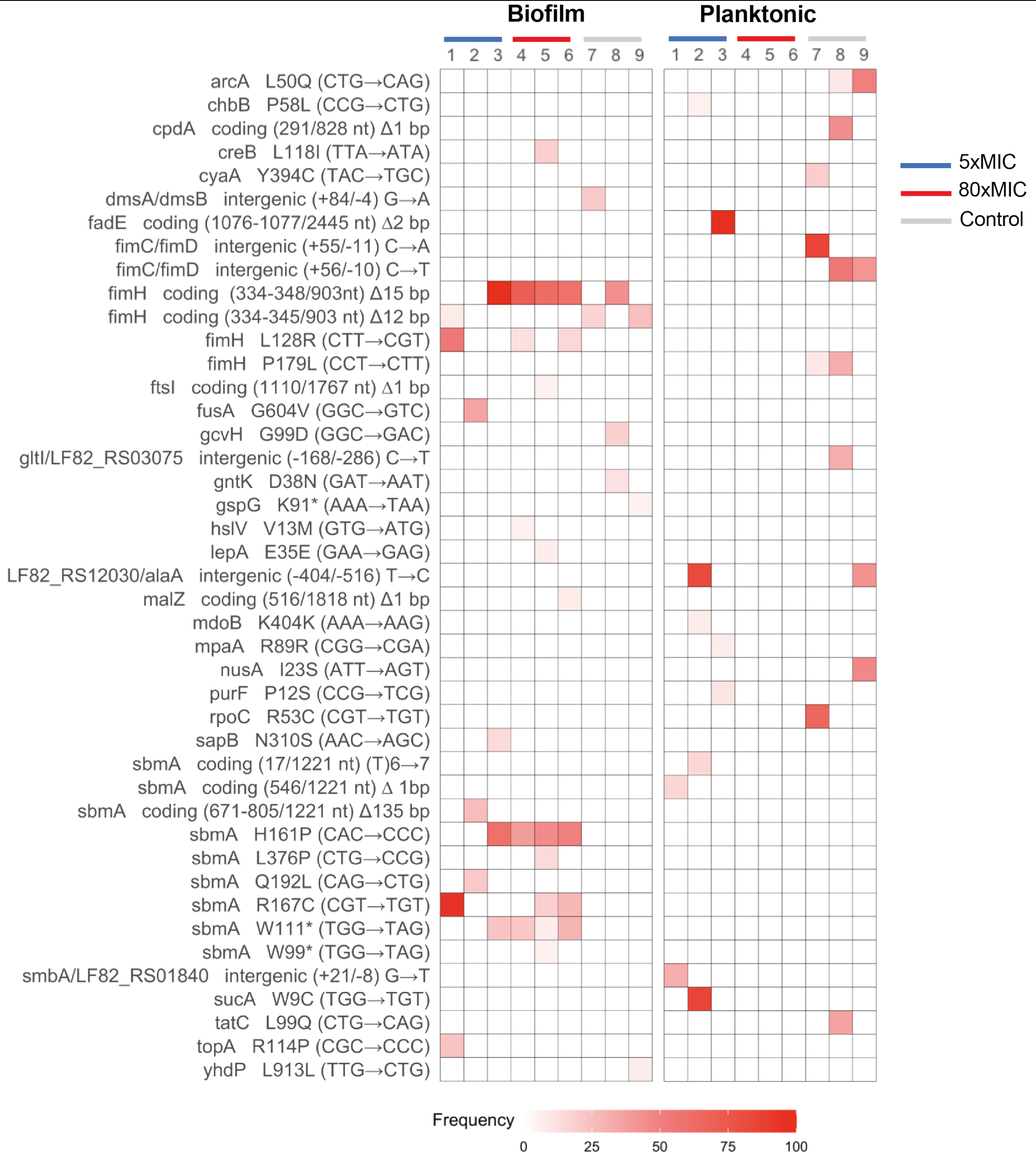


**Supplementary Fig. S6. End-point population sequencing reveals a lifestyle associated pattern of mutations after evolution under antibiotic intermittent treatment (details at the alleles level).** Mutations identified by whole-population genome sequencing of control and amikacin intermittently treated (5xMIC and 80xMIC) biofilm and planktonic populations of *E. coli*. Three populations per lifestyle and treatment were sequenced after 10 cycles of treatment. Red shading indicates the frequency of each independent mutation in the different locus at cycle 10 of the experimental evolution. Population 4, 5, 6 from the planktonic lifestyle did not survive after 3 cycles and thus could not be sequenced at cycle 10. We therefore sequenced the population corresponding to the last cycle before their respective extinction. Mutations at higher frequency than 5% are detected by Breseq analysis.

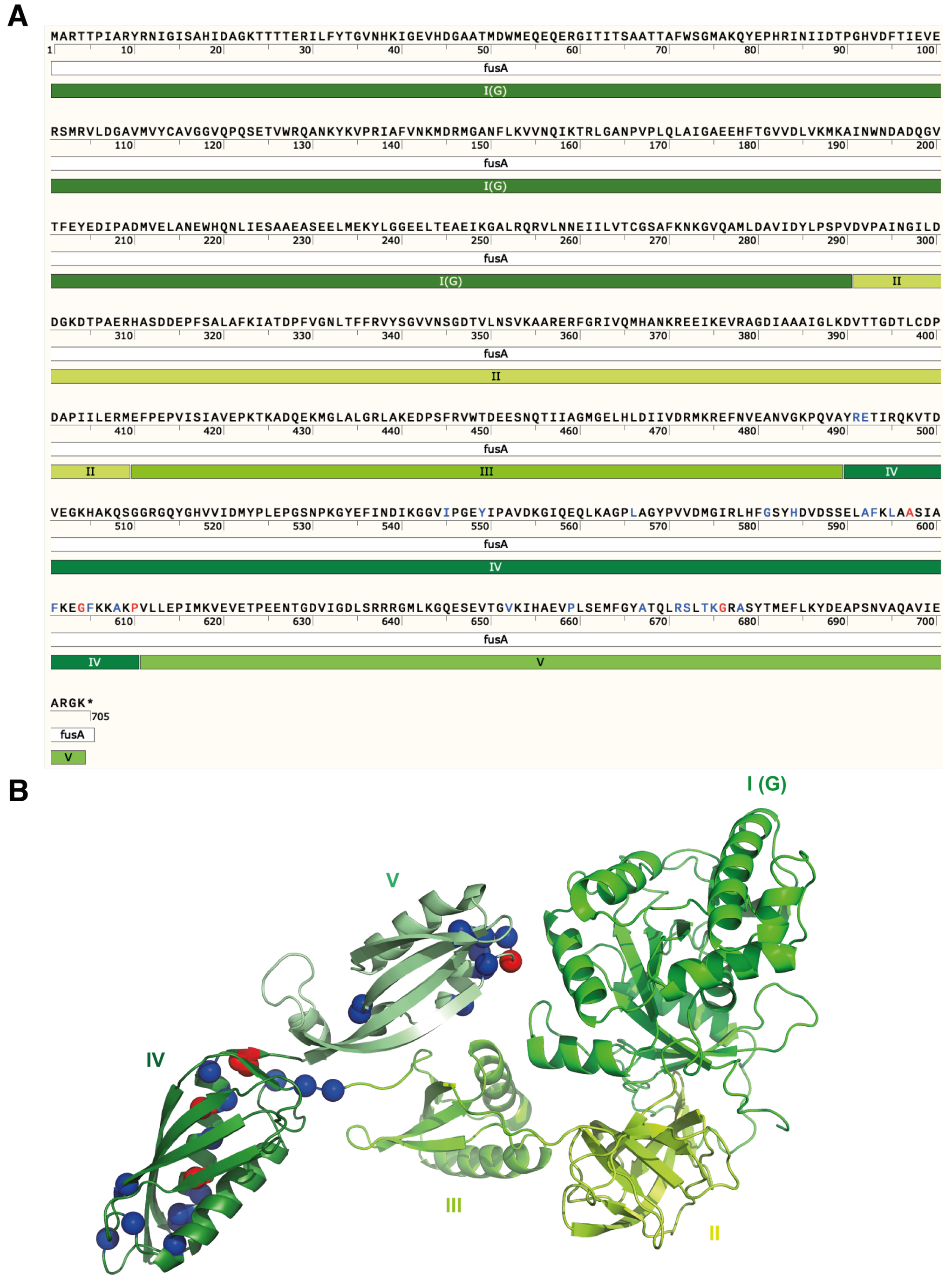


**Supplementary Fig. S7. The elongation factor G (FusA) mutations leading to amikacin resistance.** In (A) the primary sequence of *E. coli* LF82 FusA protein where the 5 functional domains are indicated. Amino-acids that correspond to positions that were identified as mutated and conferring enhanced amikacin MICs in this whole study are labeled in red for biofilm evolution on silicone coupons (A597V, G604V, P610L, G676C) and in blue for planktonic evolution (R491C, E492Q, I545T, Y549N, L566P, G581S, H584Q, A592V, F593C, F593L, L595R, F601C, F601S, F601V, F605L, A608E, A608V, P610S, P610L, V652A, P659L, P659R, A667E, R671L, S672A, T674I, K675I, A678V). Mutations in blue were detected in clones only after concentrating planktonic evolved populations. In (B), the 5 functional domains and the different mutations are represented on the model structure of LF82 FusA protein as predicted using I-Tasser (Yang & Zhang, 2015). The same color code is used that in (A). The P610 position was mutated both in planktonic and in biofilm evolution.

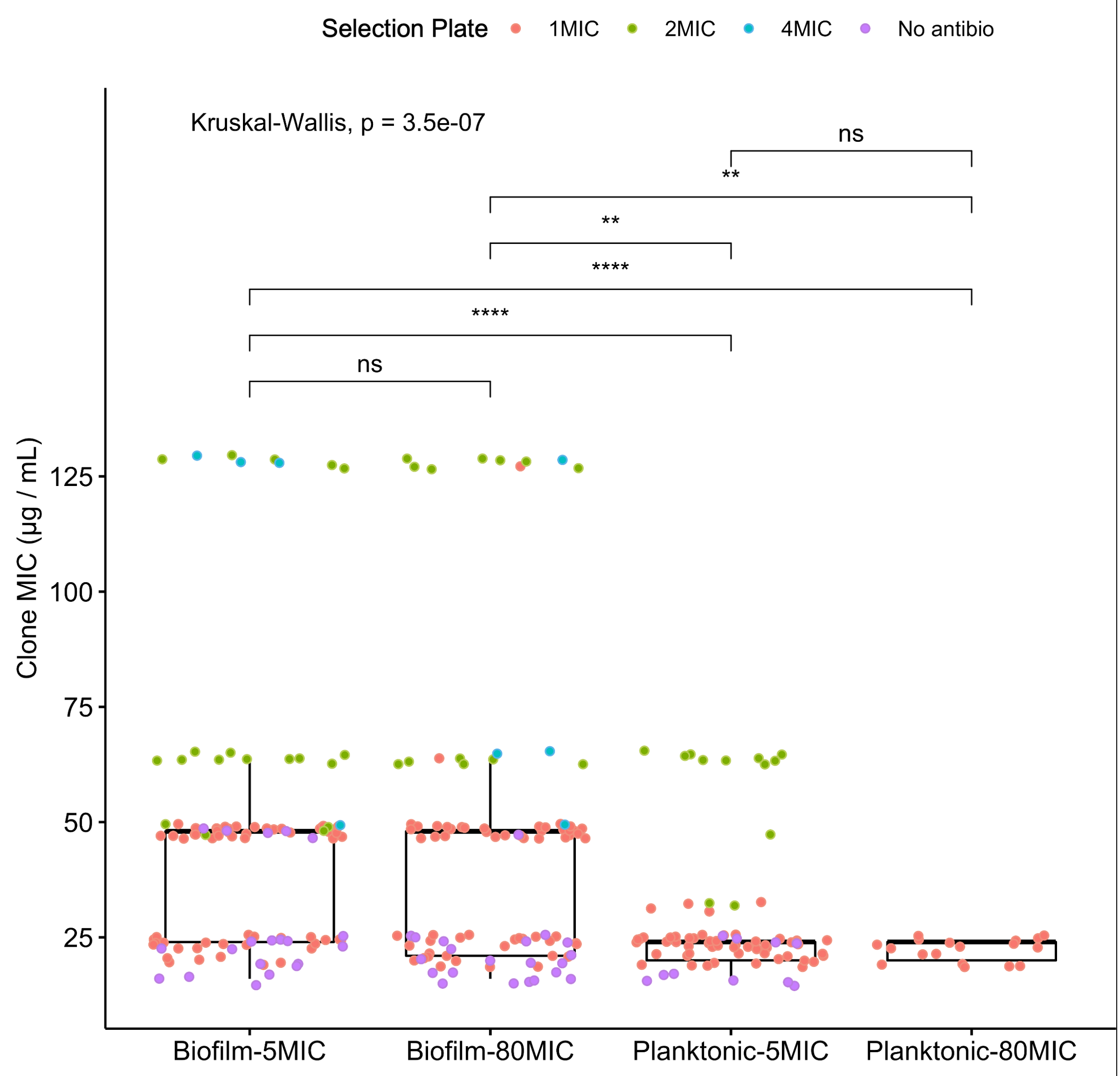


**Supplementary Fig. S8.** MIC of all clones isolated from biofilm and planktonic samples exposed to 5x and 80x amikacin MIC and summarized in supplementary Table S4, S5 and S6 (Biofilm-5MIC, n=102; Biofilm-80MIC, n=98; Planktonic-5MIC, n=74; Planktonic-80MIC, n=18)**.** For each condition the MIC for amikacin of each isolated clone is plotted, showing a higher mean MIC for the biofilm populations as compared to the planktonic populations (Kruskal-Wallis test with Benjamini-Hochberg correction). The plate on which each clone was isolated is indicated by a color code: purple on plates without antibiotic, red on plates with 1x amikacin MIC, green on plates with 2x amikacin MIC, blue on plates with 4x amikacin MIC. Statistical differences between pairs of conditions for each selection plate type were as follows (only significant ones are listed): No antibio (ns). 1MIC: Biofilm-5MIC vs Planktonic-5MIC, 6.2e-08; Biofilm-5MIC vs Planktonic-80MIC, 1.9e-05; Biofilm-80MIC vs Planktonic-5MIC, 1.2e-05; Biofilm-80MIC vs Planktonic-80MIC, 5.4e-04. 2MIC: Biofilm-80MIC vs Planktonic-5MIC, 0.005. 4MIC (ns).

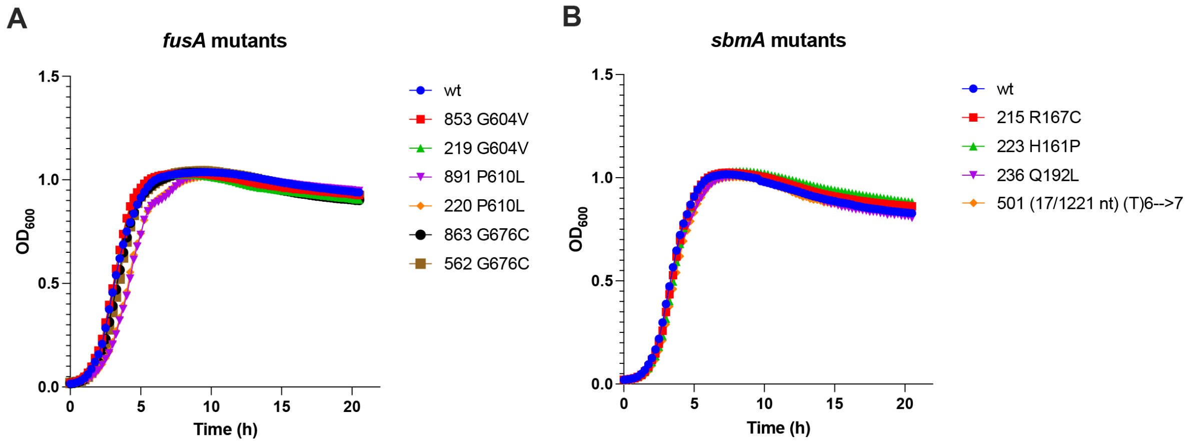


C

| | | Growth Rate ($min^{-1}$) | Generation Time (min) | Carrying capacity ($OD_{600}$) |
|---|---|---|---|---|
| | WT | 0.0256 | 27.52 | 0.962 |
| *fusA* | 853 | 0.0261 | 26.55 | 0.987 |
| | 219 | 0.0274 | 25.34 | 0.974 |
| | 891 | 0.0196 | 35.28 | 0.989 |
| | 220 | 0.0200 | 34.68 | 0.983 |
| | 863 | 0.0251 | 27.60 | 0.975 |
| | 562 | 0.0249 | 27.87 | 0.993 |
| *sbmA* | 215 | 0.0276 | 25.09 | 0.946 |
| | 223 | 0.0271 | 25.60 | 0.967 |
| | 236 | 0.0253 | 27.49 | 0.915 |
| | 501 | 0.0268 | 25.84 | 0.913 |

**Supplementary Fig. S9. Growth capacities of *fusA* and *sbmA* mutants. (A)** 20-h growth curve of *fusA* mutants in 96-well plate in LB medium. $FusA^{P610L}$ mutants (891 and 220) display slightly increased generation time compared to wt and other *fusA* mutants. (**B**) 20-h growth curve of *sbmA* mutants in 96-well plate in LB medium, showing no difference among wt and *sbmA* mutants. (**C**) Table showing the growth rate ($min^{-1}$), doubling time (min) and carrying capacities ($OD_{600}$) for all *smbA* and *fusA* mutants as inferred from the growth curves using the GrowthCurver R package (Sprouffske & Wagner, 2016).

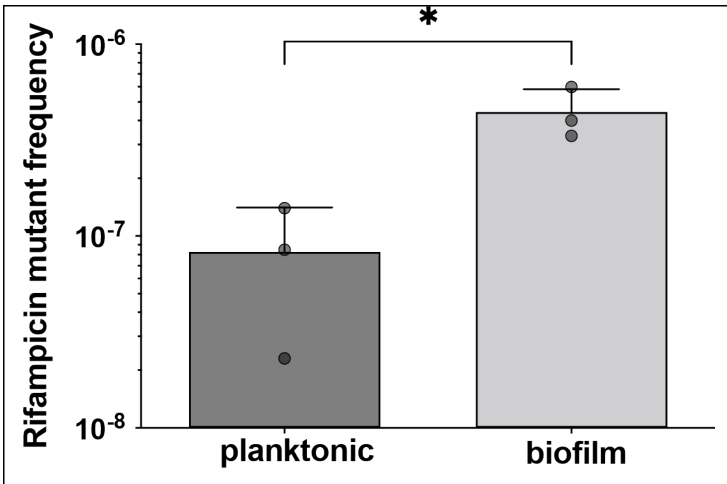


**Supplementary Fig. S10. Biofilms on silicone coupons display higher mutation rate than their planktonic counterpart.** Biofilms of LF82 strain were grown on silicone coupon during 24h at 37°C under static conditions while planktonic cells were grown under shaking conditions. After 24h of growth, silicone coupons or aliquots of planktonic culture were collected and plated on LB with or without 4xMIC of rifampicin. Frequency of resistant mutants was calculated as the $CFU_{4xMIC}/CFU_{LB}$. Statistical significance was evaluated using unpaired two-tailed t-test with Welch's correction. * $P<0.05$.

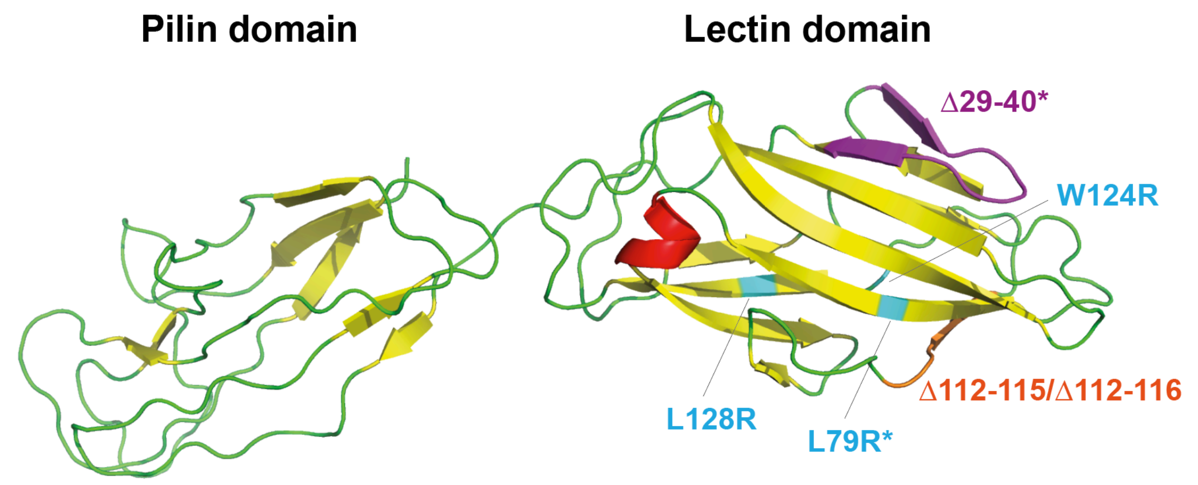


**Supplementary Fig. S11. FimH mutations selected are in the lectin domain**. Mutations identified in evolved biofilm populations are displayed on the LF82 FimH modeled 3D structure. The 3D structures of the mutant FimH proteins were predicted using the PHYRE2 Protein Fold Recognition Server (Kelley *et al*, 2015). All mutations are encompassed in the lectin domain. Mutations with a * corresponds to a deletion or a mutated position previously identified in our adaptive evolution experiments for increased adhesion (Yoshida *et al.*, 2022).

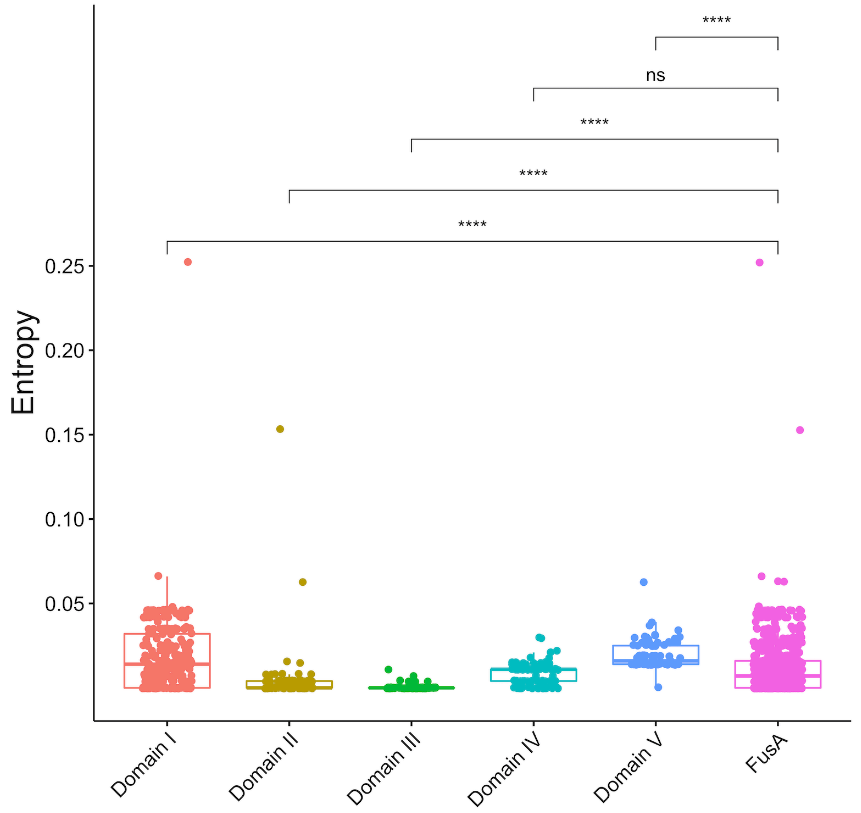


**Supplementary Fig. S12**. **Polymorphism level of different domains of the *E. coli* FusA protein.** Boxplots showing the entropy for each of the 5 domains of FusA as well as the whole FusA sequence within the multiple sequence alignment of all FusA protein sequences of *E. coli* in the nrprot database of NCBI (2143 sequences). The higher the entropy, the more variability in the alignment. Statistics correspond to one-way ANOVA followed by TukeyHSD post-hoc test comparing each domain to the whole *fusA* sequence. * p<0.05; ** p<0.01; *** p<0.001, **** p<0.0001.
